# Virulence evolution of stem and stripe rust towards the widely-deployed *Sr31*/*Yr9* wheat resistance gene combination

**DOI:** 10.64898/2026.11.23.768733

**Authors:** Jana Sperschneider, Camilla Langlands-Perry, Parastoo Karimian, David Lewis, Jibril Lubega, Eva C. Henningsen, Geetha Perera, Jian Chen, Tim C. Hewitt, Feng Li, Dhara Bhatt, Zhouyang Su, Mihir Singh, Brenton Brooks, Ming Luo, Michael Ayliffe, Brian Steffenson, Kostya Kanyuka, Rohit Mago, Melania Figueroa, Peter N. Dodds

## Abstract

The wheat *Sr31* resistance gene encoded on the 1BL.1RS wheat-rye recombinant chromosome provided durable, broad-spectrum protection against stem rust, caused by *Puccinia graminis* f. sp. *tritici* (*Pgt*), until the emergence of the virulent Ug99 lineage in 1998. Using mutant sequencing, we identify both the stem rust avirulence effector *AvrSr31* and its cognate wheat resistance gene *Sr31*. The Ug99 epidemic strain arose through a single mutation of the *AvrSr31-01* allele in a progenitor lineage that was heterozygous at this locus. The *Sr31* paralog *Yr9*, also encoded on 1RS, recognizes an unrelated effector, *AvrYr9*, from *P. striiformis* f. sp. *tritici* (*Pst*), identified here via a high-throughput effector library screen. In *Pst*, virulence on *Yr9* evolved through amino acid substitutions at two adjacent residues and expression polymorphisms that suppress AvrYr9 recognition. AvrSr31 variants in *Pgt* are more variable, with 22 polymorphic amino acid sites, but most variants are recognised by *Sr31*, except AvrSr31-03 in which two unique amino changes cause loss of recognition. These findings reveal how *Pgt* and *Pst* independently overcame two of the most widely deployed resistance genes in wheat and provide a foundation for the informed deployment of resistance against these globally important pathogens.

## Introduction

Wheat (*Triticum aestivum*) is one of the most important staple food crops worldwide. Currently, at least 8.5% of the world’s wheat production is lost each year due to fungal diseases, with over 80% of the global wheat growing areas at disease risk from the four most serious fungal pathogens (*Puccinia graminis* f. sp. *tritici*, *P. striiformis* f. sp. *tritici*, *P. triticina* and *Fusarium graminearum*) (Chai *et al*., 2022). Whilst major resistance genes bred into new plant varieties provide effective resistance, fungal pathogens can evolve quickly to overcome resistance and cause epidemics. Most resistance genes encode immune receptor proteins of the nucleotide-binding leucine-rich repeat (NLR) family, which recognise specific effectors, known as avirulence (Avr) proteins (Lo Presti *et al*., 2015; Jones *et al*., 2024; Dodds *et al*., 2024). Thus, virulence evolution in fungal pathogens is driven by mutations or deletions in effector genes, with single gene mutations sufficient to escape recognition.

Rye (*Secale cereale*) is an important source for improvement of wheat genetics through introduction of genes for improved disease resistance and abiotic stress tolerance (Crespo-Herrera *et al*., 2017). For instance, the 1BL.1RS wheat-rye recombinant chromosome contains the short arm of rye chromosome 1 translocated onto the long arm of the wheat chromosome 1B and resulted from a Robertsonian translocation event isolated during a breeding experiment in Germany in the 1930s, involving a cross between the rye cv. Petkus and wheat (Zeller & Hsam, 1973). This 1RS rye chromosome segment carries the tightly linked genes *Lr26*, *Sr31*, *Yr9* and *Pm8* conferring race-specific resistance to leaf rust (caused by *P. triticina*), stem rust (caused by *P. graminis* f. sp. *tritici*), stripe rust (caused by *P. striiformis* f. sp. *tritici*) and powdery mildew (caused by *Blumeria graminis* f. sp. *tritici*), respectively. Due to these exceptional disease resistance and favourable yield traits, wheat lines carrying the 1BL.1RS translocation were widely deployed since the 1960s and accounted for over 60% of International Maize and Wheat Improvement Center (CIMMYT) bread wheat cultivars during the 1990s (Crespo-Herrera *et al*., 2017). However, emerging virulence to *Pm8* was detected in powdery mildew populations soon after its widespread deployment in the 1970s (Bennett, 1984). Similarly, virulence to *Yr9* emerged in the 1970s and ‘80s independently in multiple regions (Stubbs, 1985; Wellings, 2011). However, *Sr31* was exceptionally durable for over 30 years until high levels of stem rust infection were recorded in a nursery in Uganda in 1998 and attributed to an aggressive strain of wheat stem rust called Ug99 (Pretorius *et al*., 2000). The Ug99 race rendered over 90% of the wheat varieties grown worldwide susceptible and has since spread widely in Africa and the Middle East causing major epidemics in eastern Africa during the 2000’s (Singh *et al*., 2011). Rust fungi are dikaryotic and contain two haploid nuclei (Henningsen *et al*., 2025). We previously showed that the clonal lineage of *Pgt* that includes Ug99 and its derivatives contains the two nuclear haplotypes hap01 and hap03 and arose by a somatic nuclear exchange event in which hap01 was donated by a clade IX isolate (hap01/hap02) and hap03 by another unknown isolate (Li *et al*., 2019). The clade II lineage of *Pgt* present in Africa prior to the emergence of Ug99 was identified as the other likely parent donating hap03 (Guo *et al*., 2022). The earliest known isolate of the clade I lineage was collected in 1972 (Guo *et al*., 2022) and is avirulent on *Sr31*, indicating that the virulent Ug99 race emerged by a mutation in this genetic background.

The *Pm8* and *Yr9* resistance genes from the 1RS rye chromosome segment have been cloned and both encode NLR proteins. *Yr9* was cloned from a *Triticale* line by gene mapping (Wang *et al*., 2025) and in parallel, from the 1BL.1RS derivative cultivar ‘Lumai 15’ (LM15) using EMS mutagenesis (Yu *et al*., 2025a). In another 1BL.1RS derivative cultivar ‘Kenong 9204’ (KN9204) (Shi *et al*., 2022a), *Yr9* is located in a 430 kbp region containing a cluster of related NLR genes (8 NLRs annotated in Shi *et al*., 2022a, 6 additional NLRs manually annotated in Yu *et al*., 2025a). *Yr9* is a member of the *MLA* family of homoeologous genes that includes the *Sr33* and *Sr50* stem rust resistance genes in wheat and rye as well as the *Mla* powdery mildew resistance genes in barley (Periyannan *et al*., 2013; Mago *et al*., 2015; Saur *et al*., 2019). *Pm8* is unrelated to this family and can be separated from *Sr31* and *Yr9* by recombination (Hurni *et al*., 2013). However, *Sr31* shows very close genetic linkage to *Yr9* (Mago *et al*., 2005) and may therefore be part of the same cluster.

Identifying Avr effectors enables understanding of pathogen evolution and informs deployment or engineering of genes for durable crop protection. For the 1BL.1RS resistance genes, only the *B. graminis* f. sp. *tritici* effector AvrPm8 has been identified so far (Kunz *et al*., 2023), and analysis of genetic variation in this gene has helped to explain the widespread pathogen virulence on *Pm8*. Extensive mutations affecting the *AvrPm8* gene likely reflect the high selection pressure exerted by the wide deployment of 1BL.1RS in wheat cultivars. However, the prevalence of an ancient gain-of-virulence mutation in *AvrPm8* was also detected and this likely preceded the *Pm8* introgression into wheat (Kunz *et al*., 2023). Here, we identify *AvrSr31* and *AvrYr9* from *P. graminis* f. sp. *tritici* (*Pgt*) and *P. striiformis* f. sp. *tritici* (*Pst*), respectively, as well as *Sr31* from wheat, and uncover the mechanisms behind the evolution of stem and stripe rust to overcome the most widely deployed resistance genes worldwide.

## Results

### Ug99 acquired virulence for *Sr31* through deletion of *AvrSr31*

Phylogenetic analysis of whole genome sequence data from ten *P. graminis* f. sp. *tritici* (*Pgt*) isolates from the Clade I clonal lineage including Ug99 revealed a clear branch point between those virulent or avirulent on *Sr31*, consistent with a single mutation event giving rise to the monophyletic group of *Sr31*-virulent strains (**Figure 1A, Supplementary Figure S1**). To identify candidate genes for *AvrSr31*, we first generated a nuclear-phased genome reference for the avirulent strain UVPgt55 (**Supplementary Table S1**) and aligned this with the Ug99 genome (Spanner *et al*., 2026). The UVPgt55 and Ug99 chromosome sequences are >99.9% identical with only ∼5,900 SNPs between them, except for a 0.9 Mbp deletion of one end of chromosome 15 in hap01 of UVPgt55 (**Supplementary Table S2**, **Supplementary Figure S2**). Because Pgt21-0 shares the hap01 nuclear haplotype with the Ug99 lineage (Li *et al*., 2019) (**Supplementary Figure S1**) and is avirulent on *Sr31*, we used existing RNA-seq data from different tissue types and Iso-seq data from infected leaves (Upadhyaya *et al*., 2015; Sperschneider *et al*., 2026) to re-annotate effector genes in the Pgt21-0 genome reference. This resulted in 8,206 genes and isoforms encoding secreted proteins and these were mapped onto the Ug99 and UVPgt55 genomes. We then mapped whole genome sequence data from *Sr31*-virulent and avirulent isolates (**Supplementary Table S3**) onto the Pgt21-0 hap01 and UVPgt55 hap03 nuclear genomes and called SNPs and structural variants. Six secreted protein gene models with expression in haustoria showed a change in the encoded protein in the virulent isolates compared to the avirulent isolates (**Supplementary Table S4**). Three of these are predicted effectors and were considered potential candidates for *AvrSr31*. These three candidates were tested by expression in protoplast cells from the wheat line Federation*4/Kavkaz (Fed/Kav) containing *Sr31* and from Fielder (lacks *Sr31*). Expression of the mature proteins encoded by AvrSr31-Candidates 1 and 3 did not induced any response in either line, while AvrSr31-Candidate2 induced cell death in protoplasts from Fed/Kav but not from Fielder (**Figure 1B**), and was designated as *AvrSr31-01*. *AvrSr31-01* is a small single-exon gene (387 bps) on chromosome 2 of hap01 in Pgt21-0 and UVPgt55 and is disrupted in the *Sr31*-virulent isolates by a 395 bp deletion starting in the fourth codon and extending to 18 bp after the stop codon (**Supplementary Figure S3A**). The alternative allele (*AvrSr31-03*) in hap03 of clade I isolates UVPgt55 and Ug99 encodes a protein with eight amino acid differences from *AvrSr31-01* that shows substantially reduced recognition in protoplasts of Fed/Kav (**Figure 1B**), consistent with the virulent phenotype of Ug99 and its derivatives. In further confirmation of the *AvrSr31* gene function, recombinant *Barley stripe mosaic virus* (BSMV) expressing *AvrSr31-01* was unable to infect Fed/Kav but did infect Cadenza, a stem rust susceptible line (**Figure 1C**). BSMV expressing the *AvrSr31-03* allele was fully virulent on both Fed/Kav and Cadenza.

**Figure 1:**
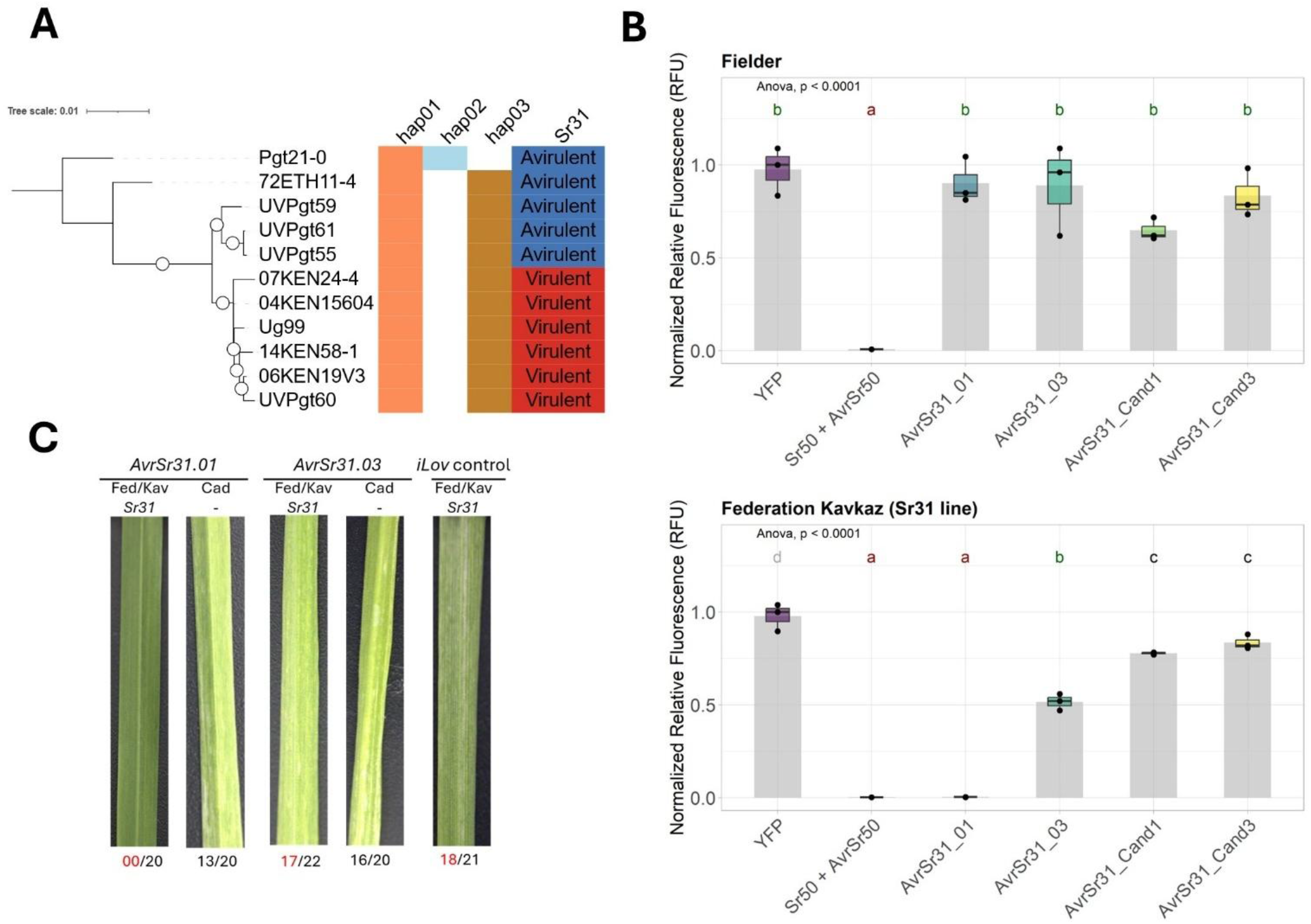
AvrSr31 identification and validation. **(A**) Maximum Likelihood (ML) tree of *Pgt* isolates based on whole genome SNPs called from Illumina sequence data aligned against the Pgt21-0 hap01 haplotype sequence. Branches with bootstrap support (100 cycles) ≥ 90% are indicated by white circles at branch midpoints. **(B)** *AvrSr31* candidate genes were expressed in protoplasts of wheat cv.s Fielder (top) and Fed/Kav (bottom) and fluorescence of a co-transformed YFP reporter gene measured (RFU, *y*-axis). Co-expression of *Sr50* and *AvrSr50* were included as a positive control. **(C)** Infection phenotypes of BSMV strains expressing AvrSr31-01, AvrSr31-03, or the small fluorescent flavoprotein iLov (negative control) after inoculation of wheat varieties Fed/Kav containing *Sr31* and Cadenza (Cad), not known to contain any stem rust resistance genes. Images show typical symptoms at 24 days post-inoculation, and the numbers below indicate the successful infections observed out of the total inoculated plants.

### Sr31 is encoded by a member of an MLA-like NLR gene cluster

To identify the *Sr31* resistance gene, we generated whole-genome sequencing data from Fed/Kav and two stem rust susceptible mutants previously isolated by mutagenesis of Fed/Kav; one that had lost only *Sr31* resistance (M2-10-1) and another that had lost both *Sr31* and *Yr9* (M2-34-5) resistance but retained all flanking markers (Mago *et al*., 2005). Sequence reads were mapped to the reference genome of Kenong9204 (Shi *et al*., 2022a), which contains the 1BL.1RS translocation carrying *Yr9* and *Sr31*. Sequencing coverage analysis showed that mutant M2-34-5 has a 2.25 Mbp deletion spanning the region from 13.47 – 15.72 Mbp on the 1BL.1RS chromosome (**Figure 2A**), which therefore delimits the location of *Sr31*. This includes a previously described 430 kbp region that harbours the cloned *Yr9* gene (15.2 – 15.65 Mbp, *Yr9* located at 15.47 Mbp) as well as a cluster of 13 related genes encoding NLR proteins, of which four are expressed (Yu *et al*., 2025b). The M2-10-1 mutant showed full sequence coverage across the 2.25 Mbp *Sr31* region and we detected only one SNP within an annotated gene; an intron 3′ acceptor splice site change from AG to AC in *NLR-6*, one of the expressed genes in the NLR cluster (Yu *et al*., 2025b). *NLR-6* is located at 15.62 Mbp, 150 kbp downstream from *Yr9*, and encodes a protein of 956 amino acids with canonical coiled-coil (CC), nucleotide-binding (NB) and leucine-rich repeat (LRR) domains (**Figure 2B**). The intron splice site mutation in M2-10-1 could result in intron retention resulting in a coding frame shift and translation of a truncated 286 aa protein (**Figure 2B**) or splicing to another potential acceptor site also disrupting the protein coding sequence. A predicted protein-protein interaction structure between AvrSr31 and NLR-6 generated with Boltz-2 (Passaro *et al*., 2025) showed a Reactifptm score of 0.951 indicating a highly confident and high-quality interface prediction. This was the highest scoring AvrSr31-interaction predicted for the 14 NLRs in this cluster (**Figure 2C**), supporting *NLR-6* as the strongest candidate for *Sr31*. In addition to the mutants described above, we generated four additional mutants by EMS mutagenesis of Fed/Kav that lost *Sr31* resistance (**Figure 2D**), but retained flanking markers and *Yr9* stripe rust resistance (**Supplementary Table S5**). The *NLR-6* gene could not be amplified from one mutant (M4-13-1), suggesting that it carries a deletion of this gene, while sequences amplified from the other three mutants contained non-synonymous mutations: a premature stop codon in M14-6-1 leading to a truncated protein of 129 amino acids and amino acid substitutions (G201R and G206S) in the highly conserved P-loop motif in mutants M14-7-1 and M14-8-1, respectively (**Figure 2C**). Lastly, we transformed the susceptible wheat cultivar Fielder with the *Sr31* candidate gene with native promoter and 3’ regulatory sequences and generated eight independent transgenic lines. All eight T0 plants showed resistance to the Pgt98 strain, which is virulent on Fielder (**Supplementary Figure S3B**). Four lines were tested again as T1 families, with two families showing segregation for the transgene and the other two uniformly resistant. In the two segregating families, the *Sr31* transgene co-segregated with resistance (**Supplementary Figure S3C**). Together, these results confirm that *NLR-6* is *Sr31*. The Sr31 protein is closely related to Yr9, Sr33 and Sr50 (pairwise amino acid identities 79-85%; **Figure 2E**), which are all orthologs of the barley *Mildew Resistance Locus A* (MLA) family of immune receptors (Bettgenhaeuser *et al*., 2021a).

**Figure 2:**
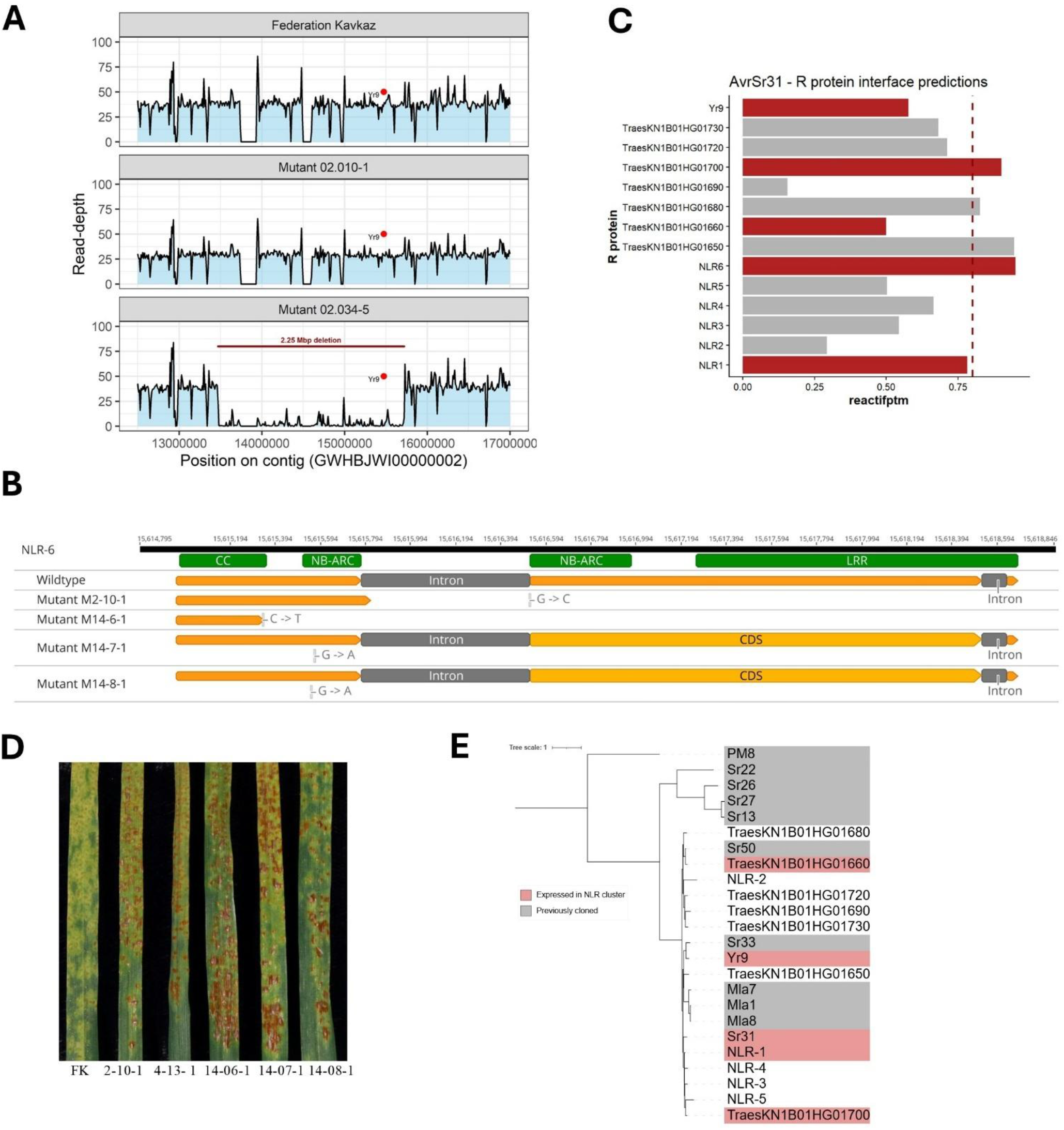
Identification of Sr31. **(A)** Sequencing coverage analysis shows that mutant M2-34-5 has a 2.25 Mb deletion (13.47 – 15.72 Mb) which includes the cloned *Yr9* gene. **(B)** Effect of mutations in the four mutants on the *NLR-6* gene. Top line shows gene structure in Fed/Kav (wildtype) with coding sequence in yellow and an intron shown in grey. Regions encoding the coiled-coil (CC), nucleotide-binding (NB)-ARC and leucine-rich repeat (LRR) domains are indicated in green. **(C)** Multimer interface confidence scores of protein-protein interaction complexes between AvrSr31 and the 14 NLR proteins in the *Yr9 -*like gene cluster. Expressed genes are shown in red. The dotted red line indicates a reactifptm score of 0.8, above which interface predictions are considered high confidence. **(D)** Infection phenotypes of Fed/Kav (FK) and five EMS-induced stem rust-susceptible mutants after inoculation with Pgt98. **(E)** A phylogenetic tree of 14 NLR proteins encoded at the *Sr31*/*Yr9* locus and additional known stem rust resistance proteins (Sr13, Sr22, Sr26, Sr27, Sr33, Sr50), MLA resistance proteins (Mla1, Mla7, Mla8) and Pm8. The 5 NLR genes at the *Sr31/Yr9* locus that are expressed (Yu et al., 2025b) are highlighted with red shading.

We further assessed *Sr31* recognition of *AvrSr31* by co-expression in wheat protoplasts, which triggered strong cell death similar to the positive control *Sr50*+*AvrSr50* (**Figure 3A**). Similarly, co-expression of *Sr31* with *AvrSr31-01* triggered cell death in *Nicotiana benthamiana* (**Figure 3B**). Four additional sequence variants of *AvrSr31* were identified in genome references of *Pgt* strains representing three other prevalent clonal lineages: Pgt21 (Sperschneider *et al*., 2026), ETH2013-1 (TKTTF) and ITA2018-1 (TTRTF) (Spanner *et al*., 2026) (**Figure 3C,D**), which are all avirulent on *Sr31*. Pgt21-0 contains *AvrSr31-01* in hap01 while the *AvrSr31-02* allele in hap02 encodes a mature protein with two amino acid differences. The hap05 haplotype of ETH2013-1 contains the *AvrSr31-05* allele, which encodes an identical mature protein to *AvrSr31-*0, although differing by four amino acids in the signal peptide region. Surprisingly, haplotypes hap04 in ETH2013-1 and hap07 in ITA2018-1 do not contain a copy of *AvrSr31* at the chromosome 2 locus, but instead contain a translocated variant, *AvrSr31-04*, in a non-allelic position on chromosome 12. The mature AvrSr31-04 protein contains eleven amino acid differences from AvrSr31-01. Hap06 of ITA2018-1 contains a 20 kbp deletion spanning the *AvrSr31* locus on chromosome 2, but no additional copy of the gene elsewhere. Expression of the AvrSr31-02/05 and AvrSr31-04 mature proteins in protoplasts of Fed/Kav triggers cell death similar to AvrSr31-01 (**Figure 3E**), indicating that they function as avirulence alleles and are recognised by *Sr31*. Similarly, BSMV strains expressing *AvrSr31-02/05* or *AvrSr31-04* are unable to infect Fed/Kav but retain virulence on wheat cv Cadenza (**Figure 3F).**

**Figure 3:**
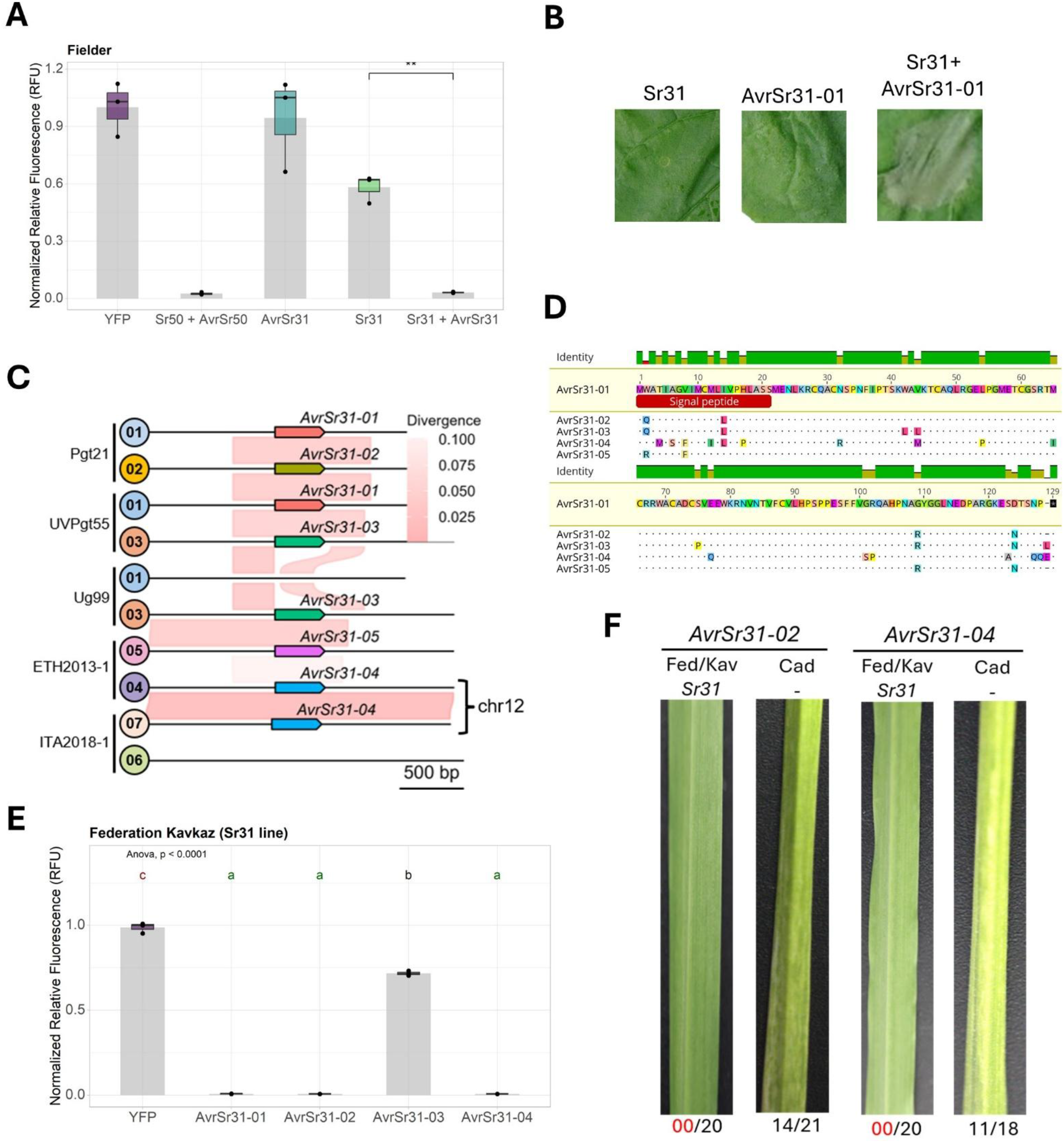
Recognition of AvrSr31 by Sr31. **(A)** *Sr31* and *AvrSr31-01* were expressed singly and in combination in wheat protoplasts (Fielder) and fluorescence of a co-transformed YFP reporter gene measured (RFU, *y*-axis). Co-expression of *Sr50* and *AvrSr50* was included as a positive control for cell death and YFP alone as a control for transformation efficiency. Reduced YFP fluorescence when R and Avr are co-expressed is indicative of immune recognition resulting in cell death. Asterisks indicate a statistically significant difference determined by a *t*-test: \*\**P* < 0.01. **(B)** *Sr31* and *AvrSr31-01* were expressed singly or in combination in *N. benthamiana* leaves by agro-infiltration. Images were recorded after 3 days to observe cell death induced by R-Avr recognition (upper panel). **(C)** *AvrSr31* locus genotypes in haplotypes from five *Pgt* isolates (Pgt21, UVPgt55, Ug99, ETH2013-1 and ITA2018-1). Gene colors represent different sequence variants with pink shading indicating sequence similarity between haplotypes. Scale for sequence divergence is shown to the right. The *AvrSr31-04* variants in hap04 and hap07 are encoded in a translocated segment on chromosome 12. *X*-axis shows the scale in bp. **(D)** Amino acid alignment of the five AvrSr31 alleles (−01 to −05). Amino acids are shown in one-letter code with dots indicating identity to AvrSr31-0 (top). **(E)** *AvrSr31-01*, *-02*, *-03* or *-04* genes were expressed in wheat protoplasts of Fed/Kav and fluorescence of a co-transformed YFP reporter gene measured (*y*-axis). Different letters on columns represent statistically significant differences (ANOVA, Tukey’s test, *P* < 0.05). **(F)** Infection phenotypes of BSMV strains expressing AvrSr31-02 or AvrSr31-04 after inoculation of wheat varieties Fed/Kav containing Sr31 and Cadenza (Cad). Images show typical symptoms at 24 days post-inoculation, and the numbers below indicate the successful infections observed out of the total inoculated plants.

### The *Sr31* paralog *Yr9* recognizes an unrelated effector from *P. striiformis* f. sp. *tritici*

To identify *AvrYr9* we used the *Yr9* gene to screen a library of effector candidate genes from *Pst* isolate Pst104 (avirulent on *Yr9*) by co-expression in wheat protoplasts (Arndell *et al*., 2024; Langlands-Perry *et al*., 2026). The effector candidates were screened in two sub-libraries of different size ranges (mature proteins less than 143 or between 143 and 363 amino acids). After library co-transformation into wheat protoplasts along with either the *Sr50* or *Yr9* resistance genes or an empty vector, we detected strong depletion of transcripts of the *AvrSr50* control in the presence of *Sr50* and of a single *Pst* effector gene (ID #874) specifically in the presence of *Yr9* (**Figure 4A, Supplementary Figure S4**). A similar library screen by transformation into wheat protoplasts from Avocet and the near-isogenic line (NIL) Avocet+Yr9 also showed reduced expression of the effector gene #874 in Avocet+Yr9 (**Supplementary Figure S4**). Co-expression of *Yr9* with #874 in wheat protoplasts from Fielder triggered cell death similarly to co-expression of the positive control genes *Sr50*+*AvrSr50*, as did expression of effector #874 alone in the Avocet+Yr9 line (**Figure 4B**). These results confirm that #874 is *AvrYr9,* and it was designated as *AvrYr9-01*. This gene encodes a protein of 262 amino acids and is located very close to the distal telomere (∼40 kbp away) of chromosome 1 (at 5.56 Mb on chr1_hap01; ∼5.34 Mb on chr1_hap02) in the Pst104 genome. The two alleles encode the same protein sequence but the hap01 allele (*AvrYr9-01-p1*) contains a 63 bp insertion in the promoter region, 228 bp upstream of the ATG start codon (∼150 bp upstream of the estimated transcription start; **Supplementary Figure S5**), relative to the hap02 allele (*AvrYr9-01-p2*). In addition to *AvrYr9*, this locus includes six distantly related effector candidate genes within a 65 kbp region (**Supplementary Figure S6**) that encode proteins that share only ∼30-45% amino acid identity with AvrYr9-01 (**Supplementary Table S6**). Three of these *AvrYr9*-homologous genes showed expression during infection and were included in the effector library but were not detected by *Yr9* in the screen (**Figure 4A, Supplementary Figure S4**), whereas the other three genes showed no transcript expression during infection.

**Figure 4:**
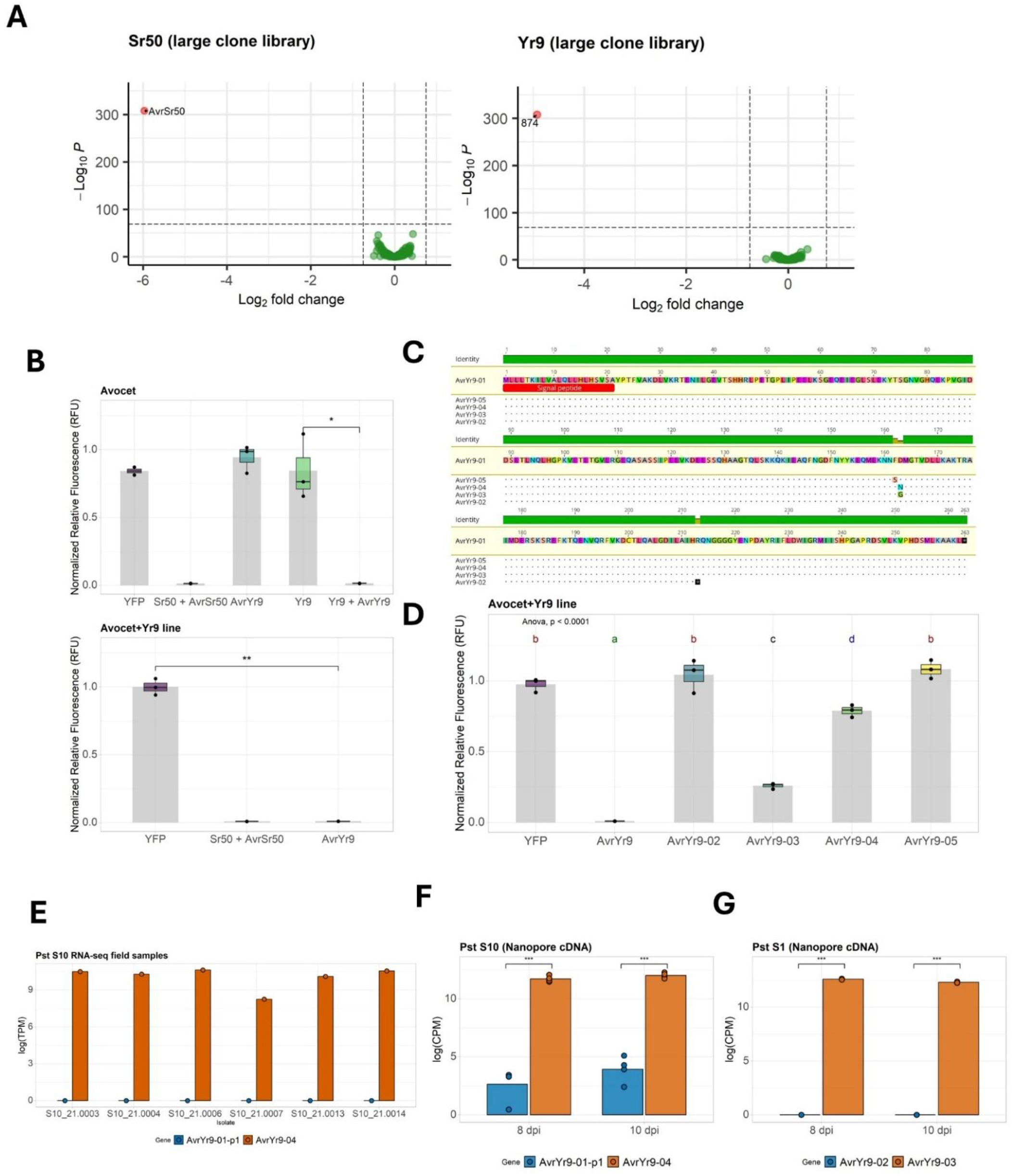
Identification of *AvrYr9* and allele-specific expression analysis. **(A)** A library of effector candidates (size range 143 to 363 aa) from *Pst* was screened against the stem rust resistance gene *Sr50* (left panel) and stripe rust resistance gene *Yr9* (right panel) by co-expression in protoplasts of wheat cv Avocet. Graphs show volcano plot of differential expression (*x*-axis) versus adjusted *p-*value (*y*-axis) for each effector construct (green dots) in the presence versus absence of the resistance gene. The positive candidate effector (clone #874) detected by *Yr9* is indicated in red as is the *AvrSr50* control. **(B)** *Sr50* and *AvrSr50* were included as a positive control for cell death and YFP alone as a control for transformation efficiency. *Yr9* and *AvrYr9-01* genes were expressed singly or in combination in protoplasts of wheat cv Avocet (top panel) and AvrYr9-01 was also expressed singly in the Avocet+Yr9 line (bottom panel) and fluorescence of a co-transformed YFP reporter gene measured (RFU, *y*-axis). *Sr50* and *AvrSr50* were included as a positive control. Asterisks indicate a statistically significant difference determined by a *t*-test: \**P* < 0.05, \*\**P* < 0.01. **(C)** A protein alignment of the AvrYr9 variants is shown, amino acids in the variants AvrYr9-02 to AvrYr9-05 that are identical to AvrYr9-01 are indicated with dots. **(D)** *AvrYr9-01*, *-02*, *-03. −04* or *-05* genes were expressed in protoplasts from Avocet+Yr9 and fluorescence of a co-transformed YFP reporter gene measured (RFU, *y*-axis). Different letters on columns represent statistically significant differences (ANOVA, Tukey’s test, *P* < 0.05). **(E)** Expression of the two alleles *AvrYr9-01-p1* and *AvrYr9-04* in RNA-seq data from six field samples of leaves infected by PstS10 strains. **(F)** Expression of the two alleles *AvrYr9-01-p1* and *AvrYr9-04* in PstS10 and **(G)** of *AvrYr9-02* and *AvrYr9-03* in PstS1 using Nanopore cDNA data.

We identified four additional allelic sequence variants (*AvrYr9-02* to *AvrYr9-05*) of *AvrYr9* in haplotype-phased genomes of 14 other *Pst* isolates, including representatives of the widespread clonal lineages PstS1, PstS7, PstS8, PstS10, PstS13 and PstS14 **(Supplementary Table S7, Supplementary Figure S7**). The *AvrYr9-02* allele has a SNP causing a premature stop, which truncates the protein by 50 amino acids, while the other three variants each have only one amino acid change relative to AvrYr9-01 (**Figure 4C**). The mature protein variants AvrYr9-02, AvrYr9-04 and AvrYr9-05 were not recognized when expressed in the Avocet+Yr9 NIL, while AvrYr9-03 was only weakly recognized (**Figure 4D**), suggesting they are virulence alleles. The *AvrYr9-02* and *AvrYr9-05* alleles contain the same 63 bp insertion in the promoter as *AvrYr9-01-p1*, while this insertion is absent from alleles *AvrYr9-03* and *AvrYr9-04* which have the *AvrYr9-01-p2* -type promoter sequence (**Supplementary Figure S7**). Notably, this promoter polymorphism correlates with the two sequence groups of *Pst* nuclear haplotypes, A and B, that are derived from two separate sub-populations in Central Asia and China, respectively (Wang *et al*., 2026b,a; Langlands-Perry *et al*., 2026). All the A-type nuclei carry alleles of *AvrYr9* with the same *p1* insertion, while the B-type nuclei contain the *p2* promoter alleles (**Supplementary Figure S8)**. In the 15 genomes, most B-type nuclei contain the *AvrYr9-04* variant (77.8%), followed by the *AvrYr9-01-p2* variant (16.7%), with the *AvrYr9-03* variant only present in PstS1. The A-type nuclei are dominated by the *AvrYr9-01-p1* (50%) and *AvrYr9-05* (33.3%) variants, with the truncation variant *AvrYr9-02* only found on hap03 in PstS14 and PstS1.

To confirm their virulence phenotypes, we inoculated Avocet and Avocet+Yr9 with Australian *Pst* isolates Pst104, 24ACT159, 22ACT160, 19NSW01 and 23NSW60, representing the PstS0, PstS1, PstS10, PstS13 and PstS14 lineages respectively. Only Pst104 was avirulent on *Yr9*, while the other four were virulent (**Supplementary Table S8**). Of these, 24ACT159 and 23NSW60 are homozygous for non-recognised *AvrYr9* alleles (−02, −03 and/or −04). However, 22ACT160 and 19NSW01 carry one copy of the *AvrYr9-01-p1* allele. Thus, we reasoned that the promoter insertion in the *AvrYr9-01-p1* allele might lead to reduced expression of this gene, to account for the virulent phenotype of these isolates. To investigate this, we used publicly available transcriptome data from other isolates of the PstS10 lineage to compare expression of the *AvrYr9-01-p1* and the *AvrYr-04* alleles, which differ by one SNP in the coding sequence allowing allele-specific gene expression analysis. The *AvrYr9-01-p1* allele had no detectable expression in RNA-seq data from field-collected infected leaves of six PstS10 isolates (Domingues Carvalho *et al*., 2026), whereas *AvrYr9-04* was expressed (**Figure 4E**). In Nanopore cDNA expression data from leaves infected by isolate Pst239 (Baig *et al*., 2026), the *AvrYr9-01-p1* variant also showed significantly lower expression than *AvrYr9-04* at time points 8 and 10 days post infection (dpi) (**Figure 4F**). We also tested allele-specific expression in Nanopore cDNA data of the isolate Pst134E (Baig *et al*., 2026), a member of the PstS1 lineage that carries the *AvrYr9-02* allele with the *p1* promoter insertion and *AvrYr9-03* without it. Again, there was no detectable expression of *AvrYr9-02* at 8 and 10 dpi, whereas *AvrYr9-03* was highly expressed (**Figure 4G**). These data confirm that the promoter insertion in the *AvrYr9-01-p1*, *AvrYr9-02* and *AvrYr9-05* alleles leads to substantially reduced expression of these genes and explains the virulent phenotype of isolates containing the *AvrYr9-01-p1* allele despite the encoded protein being identical to the *AvrYr9-01-p2* allele conferring avirulence.

### Structural modelling of AvrSr31 and AvrYr9 supports role of polymorphic residues in recognition

We previously found a number of large families of structurally-related effectors in *Pgt* and *Pst* by comparisons of AlphaFold2 predicted structures of the secretomes of these pathogens (Outram *et al*., 2026). The AvrSr31 protein is a member of an expanded structural cluster in *Pgt* that contains the zinc-binding effectors AvrSr13 and AvrSr33 as well as the AvrSr26 family (Sperschneider *et al*., 2026; Li *et al*., 2026; Outram *et al*., 2026). Similar to other members of this family, AvrSr31 has the potential to bind two zinc ions through one coordination site involving four cysteines (CCCC) and another with three cysteines and one histidine (CCCH) (**Figure 5A**). Structures predicted for the mature AvrSr31 protein by AlphaFold3 (AF3) (Abramson *et al*., 2024) support the potential for zinc-binding and places the Zn^2+^ ligands in the two expected CCCC and CCCH coordination sites (**Figure 5B**). The non-recognized AvrSr31-03 variant has two unique amino acid changes, W42L and S75P, which are exposed on the surface of the predicted structure, with the latter adjacent to one of the cysteines in the CCCC zinc-coordination site. The AF3-predicted structures for AvrSr31-01 and AvrSr31-03 showed high agreement (RMSD = 1.538) in the core structure (**Figure 5B**), but whilst S75 is predicted to be part of a beta-sheet in the recognized AvrSr31-01, the mutated residue P75 in AvrSr31-03 is part of a predicted loop structure. This localized structural distortion may contribute to the loss of recognition of AvrS31-03, or one or both of the polymorphic residues may disrupt the interaction surface with Sr31.

**Figure 5:**
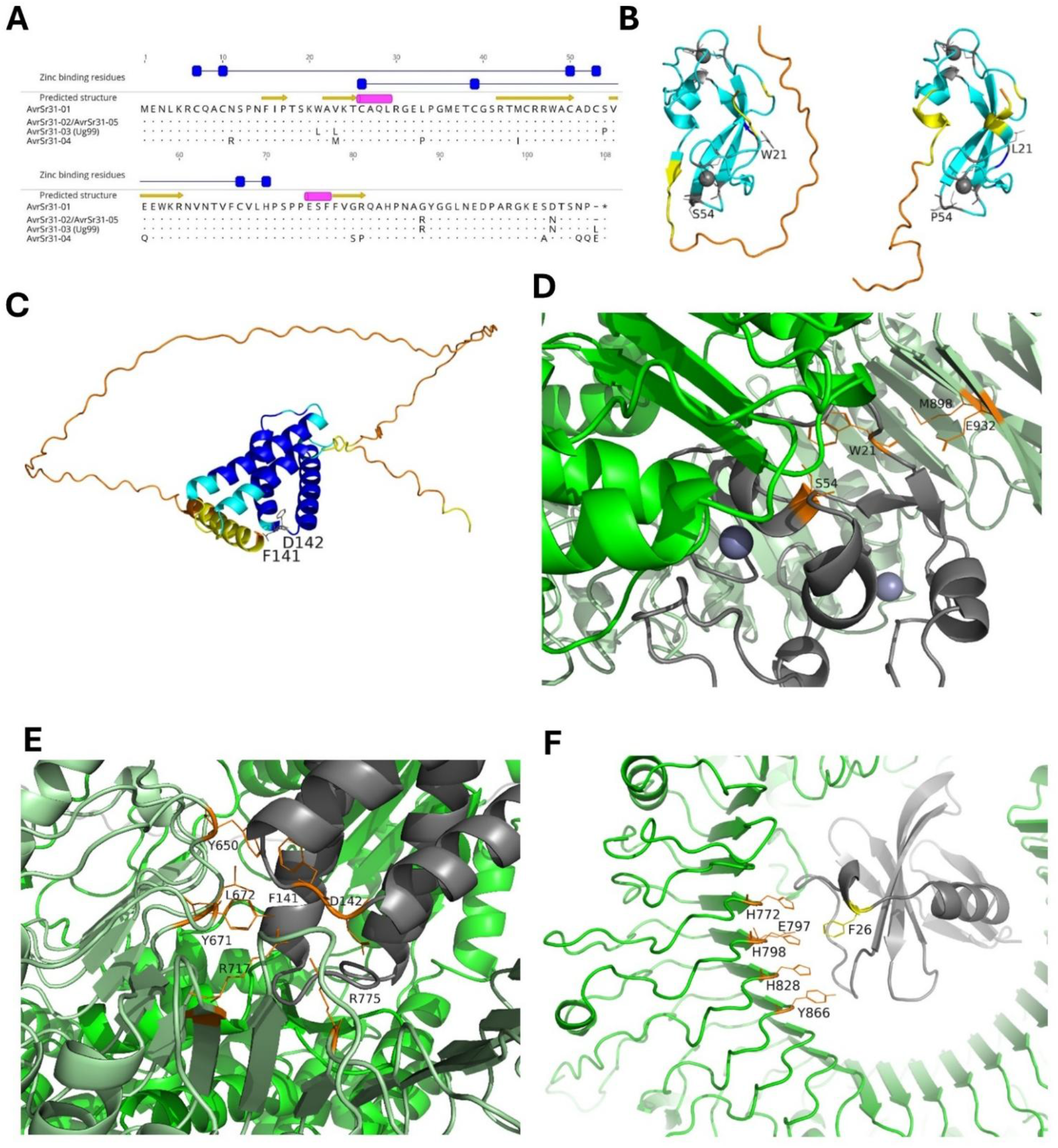
Structural predictions of AvrSr31 and AvrYr9 and recognition complexes. **(A)** Alignment of the mature AvrSr31 variant proteins. Identical amino acids are shown as dots, AF3-predicted beta-sheets are shown as yellow arrows and alpha-helices as purple boxes. Predicted zinc-binding residues are highlighted with blue, connected boxes. **(B)** The AF3-predicted structures of the mature proteins of AvrSr31-01 and AvrSr31-03 are shown on the left and right, respectively. The two mutated sites W21 (W42 in the full protein) and S54 (S75 in the full protein) are highlighted. Coloring follows the pLDDT 4-color Alphafold scheme (Blue (Very High): >90, Cyan (Confident): 70-90, Yellow (Low): 50-70, Orange (Very Low): <50). **(C)** The AF3-predicted structure of the mature protein of AvrYr9-01 is shown. The two mutated sites F141 and D142 (F162 and D163 in the full protein) are highlighted. Coloring follows the pLDDT 4-color Alphafold scheme. **(D)** AF3 predicted multimer complex between the mature AvrSr31-01 protein and Sr31 with a close-up of the region containing AvrSr31-01 residues W21 and S54. The effector is shown in gray and the R protein in green. Residues at the interface are highlighted in orange. **(E)** AF3 predicted multimer complex between the mature AvrYr9-01 protein and Yr9 with a close-up of the region containing AvrYr9-01 residues F141 and D142. The effector is shown in gray and the R protein in green. Residues at the interface are highlighted in orange. **(F)** AF3 predicted multimer complex between the mature AvrPm8 protein and Pm8 with a close-up of the region containing AvrPm8 residue F26 (F43 in the full protein sequence). The effector is shown in gray and the R protein in green. Residues at the interface are highlighted in orange.

The AvrYr9-01 protein was also included in the previous structural comparison study, but was not related to any other predicted protein structures from *Pst* (Outram *et al*., 2026). The AF3-predicted structure for the mature AvrYr9-01 protein (**Figure 5C**) contains a core region of four predicted alpha-helices with high pLDDT (>90) spanning amino acids K138 to E191 (in the full protein sequence), while the N-terminal region to amino acid L136 has low confidence (pLDDT < 50) and is largely disordered apart from a short alpha-helical region (Y22 to S43). AIUpred also predicts a disordered region spanning residues H44-T134 (probability of disorder > 0.9) in this protein. The two adjacent amino acid residues (F162 and D163) that are mutated in the non-recognized AvrYr9 variants are exposed on the surface in a predicted loop between alpha-helices in the helical bundle region.

Lastly, we predicted the complexes of AvrSr31 together with Sr31 and AvrYr9 together with Yr9. Ten residues in the Sr31 LRR domain are predicted to have close contacts (< 5 Ångström) with the mature AvrSr31 protein. Two of these, M898 and E932, are in close proximity to the W42 residue that is mutated in the non-recognised AvrSr31-03 (**Figure 5D**). For AvrYr9, the residues F162 and D163 that are mutated in the non-recognized variants are in close proximity to five Yr9 LRR domain residues (Y650, Y671, L672, R717 and R775; **Figure 5E**). We also modelled the complex of AvrPm8 and Pm8 which showed that the ancient and widespread F43Y mutation is in close proximity to five Pm8 LRR domain residues (**Figure 5F**). In summary, these structural models support the role of mutated residues in non-recognized AvrSr31 and AvrYr9 variants in Sr31 and Yr9 recognition.

## Discussion

The 1BL.1RS wheat-rye translocation carrying *Sr31* was rapidly and widely deployed in global wheats during the 1970s and later because of the value of the combined resistance traits *Pm8*, *Yr9*, *Lr26* and *Sr31*, as well as an association with increased grain yield. This provides a rare example of co-deployment of multiple disease resistance genes, allowing for comparisons of the pathogen evolutionary trajectories towards virulence in response to each gene. The *Pm8* resistance gene and corresponding effector *AvrPm8* from *B. graminis* f. sp. *tritici* (*Bgt*) were identified previously (Hurni *et al*., 2013; Kunz *et al*., 2023), while *Yr9* was identified more recently (Wang *et al*., 2025; Yu *et al*., 2025b). Here, we have identified the corresponding *AvrYr9* gene as well as the *Sr31* and *AvrSr31* interacting R-Avr pair. Although Sr31 and Yr9 are closely related, they recognise sequence- and structurally-divergent effectors. This highlights the extreme versatility of the MLA-homologous resistance gene family in Triticeae species, which has evolved a diverse array of pathogen recognition specificities. Multiple alleles of the barley *Mla* gene recognise sequence-diverse effectors from the structurally-related RNase-like effector family in the ascomycete powdery mildew pathogen *B. graminis* f.sp. *hordei* (Saur *et al*., 2019). The *Mla3* allele also recognises an unrelated effector from the blast fungus, *Pyricularia oryzae* (Gómez De La Cruz *et al*., 2026), while *Mla8* also confers resistance to wheat stripe rust (Bettgenhaeuser *et al*., 2021b) through recognition of an unknown effector. Other members of this gene family from wheat and rye, including *Sr33* (Spanner *et al*., 2026), *Sr50* (Chen *et al*., 2017)*, Sr31* and *Yr9*, all recognise structurally-unrelated effectors from basidiomycete rust pathogens.

Although deployed together as part of the 1RS resistance gene stack, the three resistance genes resulted in markedly different outcomes in terms of pathogen evolution. Both *Pm8* and *Yr9* were overcome multiple times soon after the widespread deployment of the 1BL.1RS translocation in wheat, whereas *Sr31* remained effective against stem rust for over 30 years. These contrasting outcomes are reflected in the evolutionary trajectories of their corresponding *Avr* genes. The powdery mildew effector *AvrPm8* exhibits two distinct types of mutations to overcome *Pm8* resistance: one is an ancient and globally widespread single amino acid change at F43 (to Y or L), while the 16 others are disruptive open reading frame mutations such as premature stop codons, splice-site disruptions, start-codon mutations and large-scale deletions (Kunz *et al*., 2023). The F43Y variant predates the deployment of *Pm8* and was already present in global pathogen populations, while the other 16 mutations arose recently in geographically dispersed locations and represent independent mutations to virulence. This suggests that loss of *AvrPm8* has limited fitness costs for this pathogen. The emergence and spread of virulence alleles through *Bgt* populations was likely facilitated by its haploid nature and extensive sexual reproduction (Cowger *et al*., 2016; Jigisha *et al*., 2025). In contrast, both *Pst* and *Pgt* are dikaryotic and therefore require two virulence alleles, one in each nucleus, in order to overcome resistance. These fungi are also limited to clonal reproduction through most of their geographic range due to the absence of the sexual host (*Berberis* spp). In the case of *Pst*, two separate localised sexual populations exist, one in central Asia and one in China, which contain genetically distinct genomic haplotypes designated as either A- or B-types respectively (Wang *et al*., 2026b,a). However, most globally distributed strains are hybrids between these two populations, with one A- and one B- type nucleus. Notably, the A-type nuclei in sequenced strains (AA and AB types) all contain a promoter variant of *AvrYr9* (*-p1*) that inhibits expression and results in virulence on *Yr9*, suggesting pre-existence of this virulence allele in the central Asian AA-type sexual population. Thus, strains of the PstS0 clonal lineage, such as Pst104 (isolated 1982), which were widespread globally prior to the 2000s (Wellings, 2011; Hovmøller *et al*., 2026), were already heterozygous for *AvrYr9* when *Yr9* was deployed (genotype *AvrYr9-01-p1*/*AvrYr9-01-p2*). These strains mutated to virulence on *Yr9* in multiple locations around the world likely independently (Stubbs, 1985; Wellings, 2011), which would have required only a single mutation event in the *AvrYr9-01-p2* allele. Other independent AB hybrid lineages, such as PstS1, S10, S13 and S14, have emerged this century and were all virulent on *Yr9* when first detected (Wellings, 2011; Hovmøller *et al*., 2026). These lineages contain virulence alleles in their B-type nuclei that encode single amino acid changes compared to the Avrr9-01 variant that prevent recognition. These likely arose by mutation within the sexual population in China (BB types), where virulence to *Yr9* emerged in the 1980s soon after its deployment.

In contrast to *Yr9* and *Pm8*, virulence to *Sr31* did not emerge in wheat stem rust populations until more than 30 years after its deployment. While one explanation for this could have been the presence of multiple genes on 1RS conferring stem rust resistance, here we found that a single NLR gene confers *Sr31* resistance, with mutations in this gene resulting in susceptibility. Pathogen genetics may partly explain the durability of *Sr31* resistance. For instance, the Pgt21 lineage (clade IX), which originated in Africa prior to the 1920s and was widespread globally (Pretorius *et al*., 2007; Li *et al*., 2019), is homozygous for *AvrSr31*. No mutations to virulence have been observed in this lineage. However, the clade II lineage including Ug99 was first detected in Africa in the 1970s and was already heterozygous for avirulence to *Sr31* due to the presence of the non-recognised *AvrSr31-03* allele in the hap03 nucleus. Nevertheless, it was not until 1998 that virulence to *Sr31* arose through a single mutation event deleting the *AvrSr31-01* allele in the other haplotype (hap01) of Ug99. Thus, even in a lineage requiring only a single additional mutational event to become fully virulent on *Sr31*, resistance remained effective for several decades. Historical epidemiological factors likely further contributed to this delay. Singh *et al*. (2008) proposed that the previous successful widespread deployment of other alien chromosome-derived stem rust resistance genes, including *Sr24* and *Sr26* from *Thinopyrum ponticum* and *Sr36* from *Triticum timopheevii* contributed substantially to a global decline in stem rust incidence during the 1970s and 1980s. Subsequently *Sr31* became the most extensively used stem rust resistance gene in global wheat breeding from the 1980s onwards, particularly through CIMMYT-developed cultivars. Thus, *Sr31* was initially deployed into an agricultural environment where the pathogen pressure was low and was adopted rapidly across large geographical areas, which likely caused a further decline in stem rust populations “to the extent that stem rust started to become a forgotten curse” (Singh *et al*., 2008). The low effective population size of *Pgt* during this period may have substantially delayed the emergence of *Sr31*-virulent stem rust races.

The contrasting fates of *Sr31* and *Yr9* demonstrate that extensive deployment alone does not necessarily lead to rapid resistance breakdown and highlights the importance of gene-stewardship to maintain the longevity of rust resistance genes. This includes reducing the prevalence of disease-susceptible wheat cultivars that act as reservoirs for pathogen reproduction and diversification. The deployment of *R* genes in combinations and the use of *R* genes for which the pathogen population is homozygous for the corresponding *Avr* can also serve to reduce the likelihood of virulence emergence and enhance durability by increasing the number of independent mutations required to overcome resistance. For instance, major lineages of *Pgt* are homozygous for *AvrSr22* (Spanner *et al*., 2026), while *AvrYr28* is homozygous in all sequenced *Pst* isolates (Langlands-Perry *et al*., 2026). The *Sr26* resistance gene recognises *AvrSr26* genes from multiple loci in *Pgt* (Sperschneider *et al*., 2026), equivalent to a natural resistance gene stack. It will be important to continue to identify *Avr* genes from wheat rust pathogens to inform decisions on prioritising specific *R* genes in breeding and deployment to protect wheat from these devasting diseases.

## Material and Methods

### Rust isolates and plant material

Wheat variety Federation*4/Kavkaz (Fed/Kav) containing *Sr31* on the 1BL.1RS translocation and mutant lines M02.10.-1 and M02.34-5 that have lost *Sr31* resistance were described previously (Mago *et al*., 2005). Wheat lines Fielder, Avocet and the AvocetYr9 near-isogenic line were used for protoplast expression assays while Cadenza (stem rust susceptible) was used as a control for BSMV infection. Wheat plants were grown in a growth chamber at 23°C with a 16-hour light period and 50% humidity. *Pgt* strains Pgt21, UVPtg55 and Pgt98 (race 98-1,2,(3),(5),6) have been described previously (Upadhyaya *et al*., 2015; Li *et al*., 2019). Stem rust infections were carried out as previously described (Upadhyaya *et al*., 2015). *Pst* isolates Pst104, 24ACT159, 22ACT160, 19NSW01 and 23NSW60 and infections were described previously (Langlands-Perry *et al*., 2026).

### Assembly of the UVPgt55 genome and genome comparisons

High molecular weight DNA for long-read sequencing was extracted from ∼450 mg of urediniospores as described (Li *et al*., 2019) and sequenced on the PacBio Sequel II platform at the University of Minnesota Genomics Center (UMGC). For Hi-C sequencing, 350 mg of urediniospores were cross-linked in 1% formaldehyde (20 min at ∼22 °C) and then glycine was added to a 125 mM final concentration. After 15 min incubation, urediniospores were washed twice with distilled H_2_O, centrifuged at 3500 g for 5 min, and the supernatant removed. Washed urediniospores were ground in liquid N_2_ and shipped on dry ice to Phase Genomics Inc. (Seattle, WA) for Hi-C library preparation. Paired-end libraries were sequenced with Illumina NovaSeq at Azenta Life Sciences (South Plainfield, NJ). The UVPgt55 PacBio reads were assembled with Canu 2.2.0 and default settings (Koren *et al*., 2017). The contigs were polished by aligning the PacBio reads with pbmm2 followed by one round of Arrow polishing (https://github.com/PacificBiosciences/pbbioconda). Next, the assembly was further polished with the Illumina reads and two rounds of Pilon (Walker *et al*., 2014) with the options fix --all and --mindepth 0.5 and bwa mem 0.7.17 (Li & Durbin, 2009). Contigs that have low coverage (<= 5x PacBio read coverage) were removed. Other contaminant contigs and mitochondrial contigs were removed based on Blast searches (Altschul *et al*., 1990). The assembly was phased into the two nuclear haplotypes with the NuclearPhaser pipeline which indicated that four contigs contained phase switches and had to be broken (Duan *et al*., 2022). Chromosomes were numbered according to synteny to hap01 from Pgt21-0 (Li *et al*., 2019). The haplotype chromosomes were compared to each other with mummer 4.0.0rc1, using nucmer and dnadiff to report SNPs (Marçais *et al*., 2018). Structural variants were called with SyRI (Goel *et al*., 2019) and visualized with plotsr (Goel & Schneeberger, 2022).

### *Pgt* genome annotation and variant detection

Transcripts were assembled from 6 dpi RNA-seq and Iso-Seq data and from germinated spore and haustorial RNA-seq data for Pgt21-0 as previously described (Sperschneider *et al*., 2026), merged into a consensus set with stringtie 2.2.1 (Pertea *et al*., 2015) and used as input for EffectorGeneP 1.0.0 with the *P. graminis* f. sp. *tritici* species model (Sperschneider *et al*., 2026) and the setting to use SignalP 4.1 and TMHMM 2.0 (Krogh *et al*., 2001; Petersen *et al*., 2011). RNA-seq and Iso-Seq reads were also used with Funannotate 1.8.13 (Palmer & Stajich, 2020) to run the train module (--jaccard_clip). Then CodingQuarry 2.0 was run in pathogen mode on the EffectorGeneP input transcripts (Testa *et al*., 2015) and the output provided to Funannotate predict (-ploidy 2 --optimize_augustus). Secretomes were predicted from the Funannotate annotation with SignalP 4.1/TMHMM 2.0 (Krogh *et al*., 2001; Petersen *et al*., 2011). Expression in germinated spores and haustoria was assessed with Salmon 1.10.1 (Patro *et al*., 2017) against the combined EffectorGeneP and Funannotate transcripts as well as the Pgt21-0 genome as a decoy.

Illumina genomic reads of *Sr31*-virulent isolates (Ug99, 04KEN15604, 06KEN19V3, 07KEN24-4, 14KEN58-1, UVPgt60) (Li *et al*., 2019; Guo *et al*., 2022) and of avirulent isolates (UVPgt55, UVPgt59, UVPgt61, 72ETH11-4, Pgt21-0) (Chen *et al*., 2017; Guo *et al*., 2022) were mapped to the Pgt21-0 HiFi chromosomes (Sperschneider *et al*., 2026) with bwa mem 0.7.17 (Li & Durbin, 2009). Duplicates were removed with samtools rmdup (Danecek *et al*., 2021) and SNPs were called with freebayes 1.3.7 (--use-best-n-alleles 6) (Garrison & Marth, 2012) and filtered using vcffilter of VCFlib 1.0.1 (https://github.com/vcflib/vcflib) with the parameter - f ‘QUAL > 20’. Structural variants were called with DELLY 2.6.0 (Rausch *et al*., 2012) using the same alignment files. The same process of calling SNPs and structural variants was repeated using UVPgt55 as the genome reference. The 8,206 Pgt21-0 secreted protein isoform annotation was lifted onto the Ug99 and UVPgt55 genomes with miniprot (-J20 –gff) (Li, 2023) and SNPs and structural variants were intersected with the Pgt21-0 hap01 and UVPgt55 hap03 annotations using bedtools intersect (Quinlan & Hall, 2010). Annotated gene models containing mutations were selected and candidates with expression in haustoria retained. PFAM annotations were added with InterProScan 5.72-103.0 (Jones *et al*., 2014). Effectors were predicted with EffectorP 3.0 (Sperschneider & Dodds, 2022). Avr effector protein alignments were produced with MUSCLE 5.0 (Edgar, 2004) and visualized with Geneious Prime 2026.0.2 (https://www.geneious.com).

For phylogenetic tree building. SNPs were further filtered using vcffilter of VCFlib 1.0.1 (https://github.com/vcflib/vcflib) with the parameter -f ‘QUAL > 20 & QUAL / AO > 10 & SAF > 0 & SAR > 0 & RPR > 1 & RPL > 1 & AC > 0’. Bi-allelic SNPs were selected using vcftools (–min-alleles 2 –max-alleles 2 –max-missing 0.9 –maf 0.05) (Danecek *et al*., 2011) and converted to multiple sequence alignment in PHYLIP format using the vcf2phylip script (Ortiz, 2019). Phylogenetic trees were constructed using RAxML (8.2.12) (Stamatakis, 2014). We generated 100 bootstrap trees (-f a -# 100 -m GTRCAT) and a maximum-likelihood tree (-D) and incorporated these models into a final tree (-f b -z -t -m GTRCAT) that was visualized in iTOL (Letunic & Bork, 2007).

### Transient expression in *N. benthamiana* and wheat protoplasts

The coding sequences (excluding signal peptides) of *AvrSr31* and *AvrYr9* gene variants and recombinants were codon-modified using Geneious to mimic wheat codon usage (according to this table: https://www.kazusa.or.jp/codon/cgi-bin/showcodon.cgi?species=4565) and synthesised directly in pTA22 (Twist Bioscience). Plasmids used in this study are listed in **Supplementary Table S9**. For protoplast assays, plasmids were purified using the NucleoBond Xtra Midi EF kit (Macherey-Nagel). Seedlings of wheat cultivars Fielder, Fed/Kav, Avocet and Avocet+Yr9 NIL were grown in Martins Seed Raising and Cutting Mix at 24°C under a 12 h light (∼130 µmol m-2 s-1) and 12 h dark cycle. At seven days, the first leaf was harvested for protoplast isolation and cells transformed as described (Spanner *et al*., 2026). After 20 h incubation sample aliquots were transferred in duplicate to a Black 96-Well Immuno Plate (Thermofisher, Cat. #9502867) and YFP fluorescence measured using a CLARIOstar Plus plate reader (BMG Labtech; the top optic was used, optic settings: presetname = YFP, excitation = 497-15, dichroic filter = 517.2, emission = 540-20, focal height = 5.7mm, scan settings: number of flashes per scan = 4, scan mode = matrix scan, scan matrix dimension = 30x30, scan width = 7mm). The gain was set based on fluorescence scanning of YFP control samples (scan settings as above except: number of flashes per scan = 100, scan mode = spiral average). Fluorescence was normalized against the mean of the YFP control samples for each experiment.

*Nicotiana benthamiana* and *N. tabacum* plants were grown in a growth chamber at 23 °C with a 16-hour light period and used for agroinfiltration at the 4-week-old stage. *Agrobacterium tumefaciens* cultures were grown at 28°C overnight with shaking at 200 RPM in Luria-Bertani liquid medium with appropriate antibiotic selections. Cells were pelleted by centrifugation and resuspended to a final concentration of OD_600_ = 0.2 for the *Agrobacterium* strains carrying *R* genes and OD_600_ = 0.5 for those carrying *Avr* genes in 10 mM MES pH 5.6, 10 mM MgCl_2_, 150 μM acetosyringone and incubated at 20-23.5°C for 2 h before infiltration. Leaves were photographed or scanned 3 to 5 days after infiltration.

All statistical analyses were performed using R 4.4.0. Significance of differences were evaluated using a one-way analysis of variance (ANOVA) via the aov function. Pairwise comparisons between group means were conducted using the Tukey’s Honest Significant Difference method via the TukeyHSD function. Statistical significance was defined as an adjusted *p*-value < 0.05. *T-*tests were run using the ggsignif package (Ahlmann-Eltze & Patil, 2021).

### Virus-mediated *in planta AvrSr31* effector variants expression assay

The barley stripe mosaic virus (BSMV) mediated protein overexpression system (BSMV-VOX), comprising three T-DNA binary plasmids, pCaBS-α, pCaBS-β, and pCa-γb2A-LIC, was used in this study (Kanyuka, 2022). Mature coding sequences of the *AvrSr31* variants (*AvrSr31-01*, *AvrSr31-02, AvrSr31-03, AvrSr31-04*), in which the predicted signal peptides were replaced by a methionine start codon, were PCR-amplified using primers flanking each coding sequence and carrying 5′ extensions (forward primer: CCAACCCAGGACCGTTGATG; reverse primer: AACCACCACCACCGTTA) to enable ligation-independent cloning into pCa-γb2A-LIC (primers listed in **Supplementary Table S10**). Amplified fragments were cloned into pCa-γb2A-LIC following the procedure of Kanyuka (2022), and all constructs were verified by whole plasmid sequencing. The verified BSMV binary plasmids were transformed into chemically competent *A. tumefaciens* strain GV3101(pMP90) by freeze-thaw method (Weigel & Glazebrook, 2006). Agroinfiltration of 3-4-week-old *N. benthamiana* plants proceeded, performed as described previously (Kanyuka, 2022). Crude sap from infiltrated *N. benthamiana* leaves was subsequently used to mechanically inoculate the first and second leaves of three-leaf-stage wheat plants of cultivars Fed/Kav (*Sr31*) and Cadenza (*sr31*). BSMV:iLov served as a negative control (Franco-Orozco *et al*., 2017). Inoculated wheat plants were maintained in a controlled-environment growth room at 23°C/20°C day/night temperatures, 60-65% relative humidity, and a 16 hr photoperiod with ∼220 μmol m⁻² s⁻¹ light intensity. The presence or absence of viral symptoms in systemic leaves was assessed at 10, 16, and 24 days post-inoculation. For each time point, the number of plants displaying systemic symptoms was recorded relative to the total number inoculated. Each experiment included 18-22 individual plants per BSMV-VOX construct.

### Genome analysis of *Sr31* mutants

Genomic DNA from Fed/Kav and *Sr31* mutants was extracted using the Macherey-Nagel™ NucleoSpin™ Plant II, DNA extraction kit and sequenced using the Illumina NovaSeq X Plus (300 cycles) at the Ramaciotti Centre for Genomics (Australia). Illumina reads were cleaned with fastp v.0.23.2 (Chen *et al*., 2018) and mapped to the reference genome of KN9204 (Shi *et al*., 2022a) with bwa mem 0.7.17 (Li & Durbin, 2009). Duplicates were removed with samtools rmdup (Danecek *et al*., 2021) and SNPs were called with freebayes 1.3.7 (--use-reference-allele). SNPs were filtered using vcffilter of VCFlib 1.0.1 (https://github.com/vcflib/vcflib) with the parameter -f ‘QUAL > 20’. Coverage was calculated with pileup.sh from bbmap v39.01 (https://sourceforge.net/projects/bbmap/) and binsize=10000. Structural variants were called with DELLY 2.6.0 (Rausch *et al*., 2012) using the same alignment files. The genome annotation from KN9204 (Shi *et al*., 2022a) and the additional NLR annotations (Yu *et al*., 2025b) were intersected with SNPs and structural variants to find candidates for *Sr31*.

### Isolation of additional *Sr31* mutants and candidate gene sequencing

Mutants with loss of *Sr31* resistance were generated using Ethylmethane sulfonate (EMS) treatment as described (Mago *et al*., 2017). In brief, 900 seeds of the wildtype Fed/Kav line were treated with 0.4% EMS. M2 families were screened for loss of resistance to Pgt98. Susceptible plants were recovered and retested by rust inoculation in the M3 generation to confirm the loss of resistance. Confirmed mutants were scored with DNA markers flanking the *Sr31*/*Yr9* locus (Mago *et al*., 2005) and inoculated with stripe rust isolate Pst104 to assess *Yr9* function. A 3.8 kbp region spanning the coding region of the *Sr31* gene candidate NLR6 was amplified from the confirmed mutants using the primers NLR6-F2 and NLR6-R3 and sequenced by Sanger sequencing using overlapping primers (**Supplementary Table S10**).

### Wheat transformation

Scutellum tissue from wheat cultivar Fielder was transformed with *Agrobacterium* strain AGL0 containing an *Sr31* construct (including 2.5kb upstream and 1.5kb downstream of coding sequence) using BAR (30μg/mL) for selection as previously described (Richardson *et al*., 2014). T_0_ lines were infected with Pgt98 and rust reactions were assessed after 10-15 days. T_1_ seed was subsequently harvested, and progeny of each line were grown in a growth cabinet (23°C, 16 h light). Ten-day-old seedlings were inoculated with Pgt98 and rust reactions were assessed after 10-15 days. Plants were screened for the presence of the transgene by PCR with primers NLR6-F3 and NLR6-R4.

### Effector library screening in wheat protoplasts

A library of *Pst* effector candidates in two sub-libraries of small and large size ranges (25-142 and 143-363 amino acids, respectively) was described previously (Langlands-Perry *et al*., 2026). Libraries were propagated in ElectroMAX™ Stbl4™ competent cells and DNA prepared using the ZYMO Research ZymoPURE II Plasmid Midiprep Kit. The empty vector (pTA22), pTA22-Yr9 and pTA22-Sr50 plasmids were prepared from *E. coli* DH5α using the Macherey Nagel NucleoBond Xtra Maxi Plus EF kit. For library screening a total of 300,000 protoplast cells of wheat cv Avocet were transformed with an effector library at a multiplicity of transfection (MOT) of 0.14 million molecules per cell for each construct along with either pTA22, pTA22-Yr9 or pTA22-Sr50 (MOT = 36 million molecules per cell) in triplicate as described (Arndell *et al*., 2024). The large size effector library was also transformed in triplicate into wheat lines Avocet and Avocet+Yr9 (MOT = 0.14 million molecules per cell for each construct) with no other plasmids. Total RNA was extracted from transformed protoplasts after 20 hours using the Maxwell® RSC Plant RNA Kit (Cat. #AS1500) with the Maxwell® RSC 48 Instrument (Cat. #AS8500). Library-specific cDNA synthesis and PCR was carried out on 1µg RNA using the Invitrogen SuperScript IV One-Step RT-PCR System with ezDNase kit (Cat. # 12595100) with the forward and reverse primers ZmUbi1_5UTR_F3b (5’-GCACACACACACAACCAG-3’) and FS_cDNA_R (5’-TGCTAGATCTCGACAGTACG-3’) as previously described (Arndell *et al*., 2024). The cDNA was sequenced at AGRF (Melbourne, Australia) on the NovaSeq X with 150 bp paired-end reads. Sequencing reads were cleaned using fastp 0.23.2 (Chen *et al*., 2018) and aligned to the coding sequences of the effector candidates with the vector-derived UTRs added using STAR 2.7.9a (Dobin *et al*., 2013) (--alignIntronMax 1). Read counts were extracted with samtools idxstats (Danecek *et al*., 2021) and differential expression analysis was carried out using DESeq2 (Love *et al*., 2014) with default parameters followed by lfcShrink (type="apeglm"). Volcano plots were produced with EnhancedVolcano (https://github.com/kevinblighe/EnhancedVolcano).

### *AvrSr31* and *AvrYr9* allelic variant identification

The annotation of *AvrSr31-01* from hap01 (Pgt21-0) and *AvrYr9-01* from hap01 (Pst104) were used as the reference input for liftoff v1.6.3 (-copies -flank 0.25 -sc 0.8) (Shumate & Salzberg, 2021) with other haplotypes used as targets. Gffread 0.12.7 (Pertea & Pertea, 2020) was used to correct the resulting gff files prior to extraction of CDS and protein sequences with agat 1.0.0 (Dainat *et al*., 2022). Haplotypes with no alleles reported were aligned to the reference sequence with Clustal-Omega 1.2.2 (Sievers *et al*., 2011) and/or progressiveMauve 2.4.0 (Darling *et al*., 2004) and syntenic regions were inspected in Geneious Prime 2026.1.3 (https://www.geneious.com) to characterize knockout variants. Synteny plots were generated by obtaining pairwise alignments between specified intervals using minimap2 2.25 (Li, 2018) and visualizing alignments and annotations using gggenomes 1.2.0 (Hackl *et al*., 2024). For *Pgt*, we used the following genome assemblies for variant identification: UVPgt55 (assembled here); Pgt21-0 (Sperschneider *et al*., 2026); ETH2013, ITA2018and Ug99 (Spanner *et al*., 2026). For *Pst*, we used the following genome assemblies for variant identification: Pst104, 19NSW01, 22ACT160, 23NSW60, 24ACT159 (Langlands-Perry *et al*., 2026); PstS7 (Hovmøller *et al*., 2026); AZ2 (Wang *et al*., 2024); GSY16-9, GSY17-10, A153, XZ2, SC13-4, Pak-1, Eth-1 (Wang *et al*., 2026a); DK02d-12 (Mojerlou *et al*., 2026). Genome accession numbers are provided in **Supplementary Table S7**.

### *AvrYr9* allele-specific expression analysis

RNA sequencing reads from 17 phenotyped PstS10 field samples were downloaded (S10_21.0001, S10_21.0003, S10_21.0004, S10_21.0005, S10_21.0006, S10_21.0007, S10_21.0008, S10_21.0009, S10_21.0010, S10_21.0013, S10_21.0014, S10_21.0015, S10_21.0016, S10_21.0018, S10_21.0019, S10_21.0020, S10_21.0021; NCBI BioProject: PRJNA1231252) (Domingues Carvalho *et al*., 2026). These reads were cleaned with fastp v.0.23.2 (Chen *et al*., 2018) and then mapped to the *Pst* 22ACT160 isolate (S10) genome (Langlands-Perry *et al*., 2026) with STAR 2.7.9a in 2-pass mode (--alignIntronMin 5 --alignIntronMax 3000 --alignMatesGapMax 3000 --outFilterMultimapNmax 100) (Dobin *et al*., 2013). For allele-specific expression, each sample was quantified with Salmon 1.10.1 (Patro *et al*., 2017) against the combined AvrYr9 and AvrYr9 homolog transcripts as well as the genome as a decoy. Only samples with at least five reads aligned against *AvrYr9* were used for allele-specific expression analysis. For the Nanopore cDNA data (Baig *et al*., 2026) (NCBI BioProject: PRJNA1457771), we used pychopper 2.7.10 to clean the reads (https://github.com/epi2me-labs/pychopper). Clean reads were aligned to the AvrYr9 transcripts with minimap2 (--eqx -N 100 -ax map-ont) (Li, 2018) and expression was quantified with oarfish (--filter-group no-filters -- model-coverage) (Jousheghani & Patro, 2024).

For estimating the transcription start site for the *AvrYr9* gene, we mapped RNA-seq reads of haustoria and infected leaves (6 dpi) (Schwessinger *et al*., 2018) with STAR 2.7.9a in 2-pass mode (--alignIntronMin 5 -- alignIntronMax 3000 --alignMatesGapMax 3000 --outFilterMultimapNmax 100) (Dobin *et al*., 2013) and PacBio Iso-Seq reads (Langlands-Perry *et al*., 2026) with minimap2 (-ax splice:hq -uf -G 3000) (Li, 2018) to the Pst104 genome and visually inspected the UTR regions in IGV (Robinson *et al*., 2011).

### Structure predictions

Protein complexes between the mature AvrSr31 protein and the NLR proteins were predicted with Boltz-2 (2.2.0) (-- use_potentials --recycling_steps 5 --diffusion_samples 10) (Passaro *et al*., 2025), providing a multiple sequence alignment file for each protein generated by MMseqs2 (17-b804f) (Steinegger & Söding, 2017) against UniRef90 (downloaded on 23-09-2025). Interface confidence scores were calculated with reactifpTM (Simpkin *et al*., 2026). A phylogenetic tree from resistance protein sequences was calculated using MUSCLE 5.0 (Edgar, 2004) and IQ-TREE 2 (Minh *et al*., 2020). We used AlphaFold v3.0.0 with default parameters to predict effector structures and effector-R complex structures (Abramson *et al*., 2024). For AvrSr31, we added two zinc ions to the Alphafold json input file. Structures were visualized with PyMOL (Schrödinger, LLC, 2015). We used AIUpred for disorder predictions (Erdős & Dosztányi, 2024).

## Data availability

The UVPgt55 PacBio and Hi-C reads and genome assembly and the *Sr31* mutant genomic reads are available at the CSIRO Data Access Portal at https://data.csiro.au/collection/csiro:79153v1.

## Acknowledgements

This project was supported by the Grains Research and Development Corporation project CSP2403-014RTX, the 2Blades Foundation and the Gatsby Foundation, BBSRC grant BB/W018403/1 as part of the NSF/BBSRC Lead Agency Agreement, the UK Agriculture and Horticulture Development Board UKCPVS project 21120034a, the CSIRO Research Office, and the Lieberman-Okinow Endowment at the University of Minnesota. We acknowledge Narayana Upadhyaya and Xioadi Xia for discussions and assistance. We thank the CSIRO High-Performance Computing Services for computational resources.

## Supplementary Figures

**Supplementary Figure S1:**
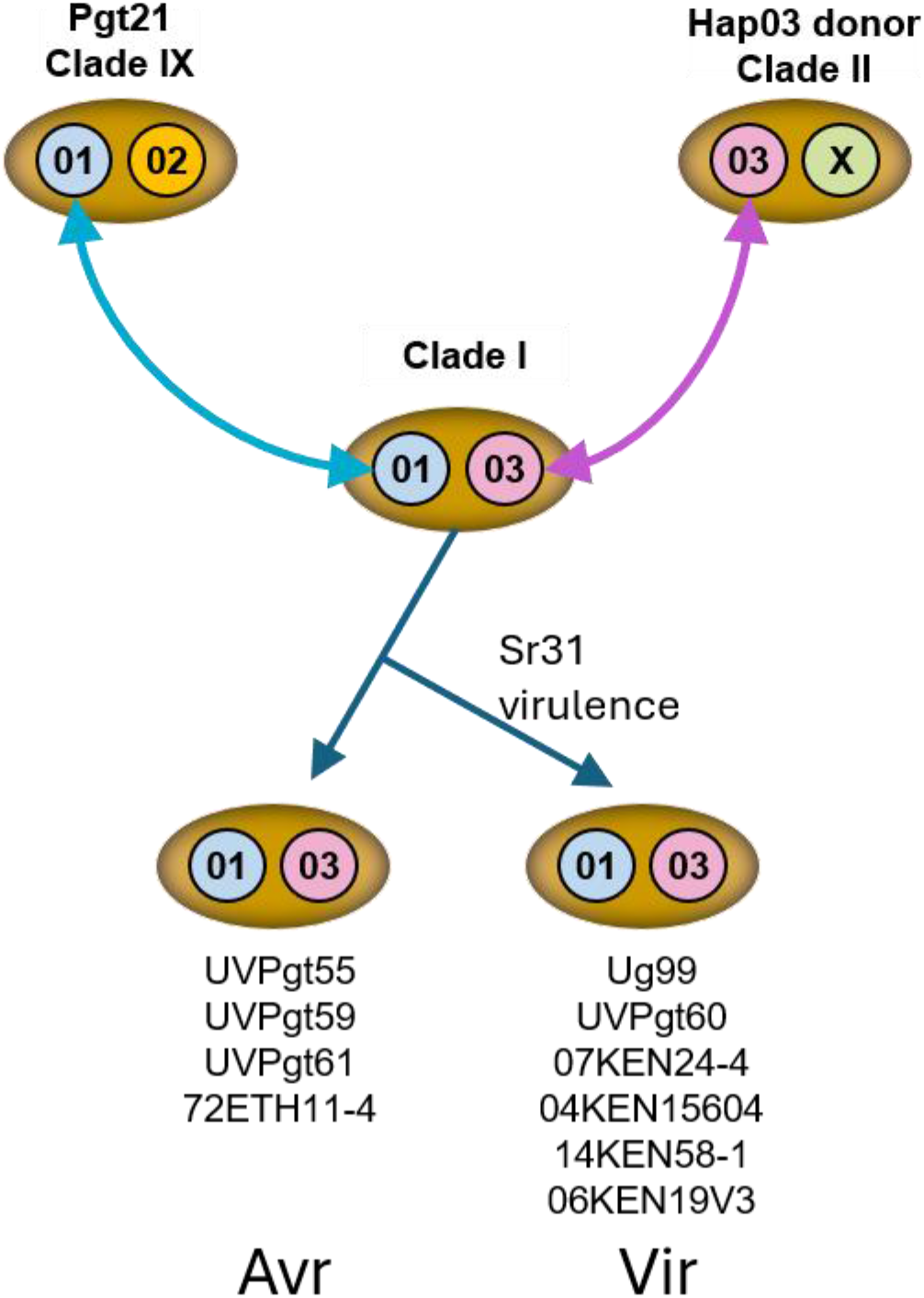
A diagram showing origins of Ug99 clade I by somatic hybridisation and nuclear exchange between Clade IX and clade II nuclear donors and subsequent evolution of virulence on *Sr31*.

**Supplementary Figure S2:**
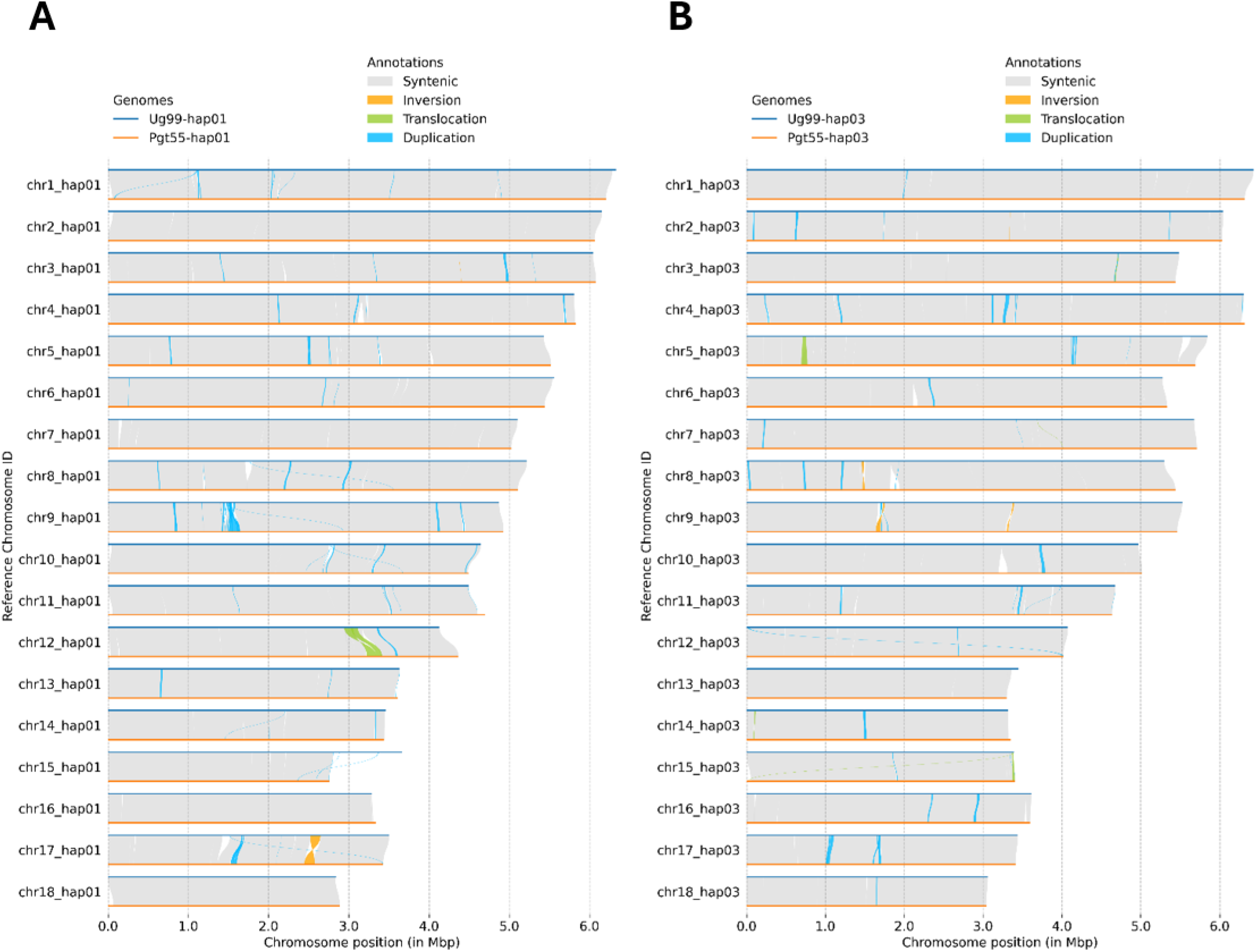
Synteny plots between the Ug99 and UVPgt55 chromosomes in haplotypes hap01 (**A**) and hap03 (**B**).

**Supplementary Figure S3:**
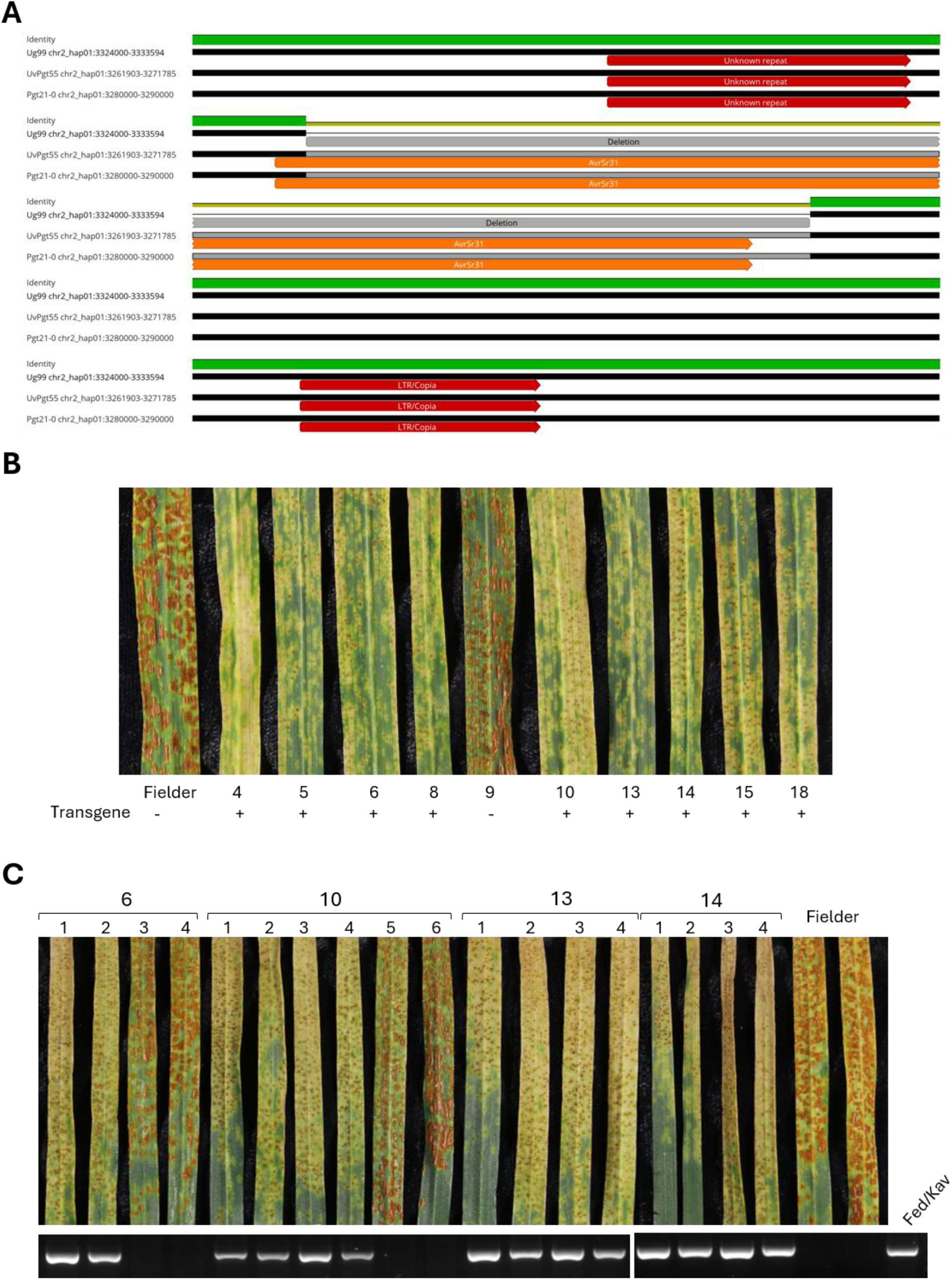
(**A**) Deletion of the *AvrSr31* gene on Ug99 hap01. (**B**) Infection phenotypes of wheat cv Fielder and T0 transgenic lines (4, 5, 6 etc) produced by transformation of Fielder with the *Sr31* candidate gene after inoculation with Pgt98. Presence or absence (+/-) of the transgene in each line as detected by PCR is indicated. (**C**) Infection phenotypes of wheat cv Fielder and four to six plants from four T_1_ families derived from T0 plants #6, #10, #13 and #14 after inoculation with Pgt98. Lower panel shows products generated by PCR amplification of each plant with the gene specific primers NLR6-F3/R4.

**Supplementary Figure S4:**
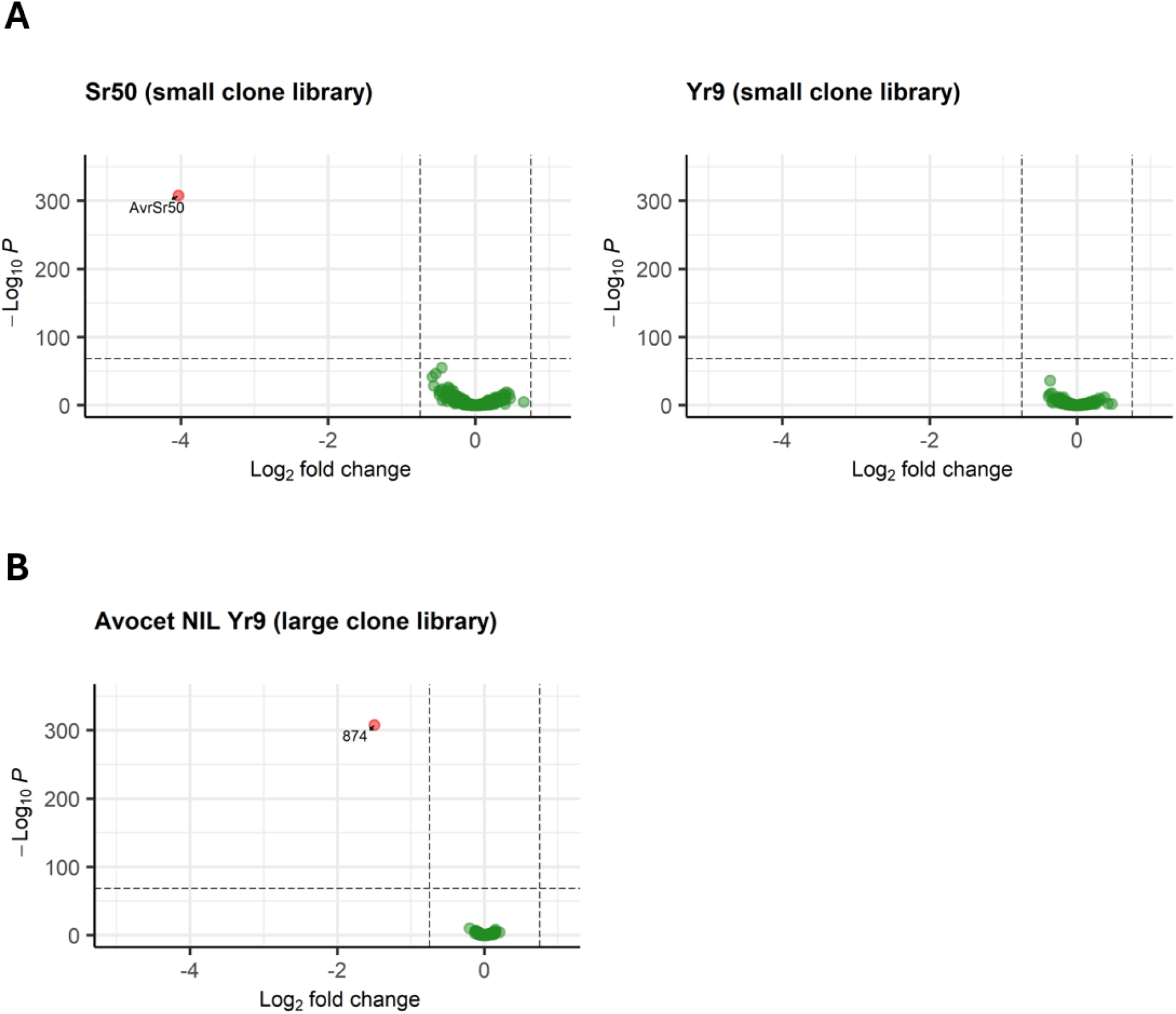
(A) A library of small (<143 aa) effector candidates from *Pst* was screened against the stem rust resistance gene *Sr50* (left panel) and stripe rust resistance gene *Yr9* (right panel) by co-expression in protoplasts of wheat cv Avocet. Graphs show volcano plot of differential expression (*x*-axis) versus adjusted *p-*value (*y*-axis) for each effector construct (green dots) in the presence versus absence of the resistance gene. (**B**) A library of larger (143 to 363 aa) effector candidates was expressed with no co-transformed *R* gene in the Avocet+Yr9 NIL line and compared to expression in the near-isogenic *Pst*-susceptible line Avocet. The positive candidate effector (clone #874) detected in the *Yr9* screens is indicated in red as is the *AvrSr50* control.

**Supplementary Figure S5:**
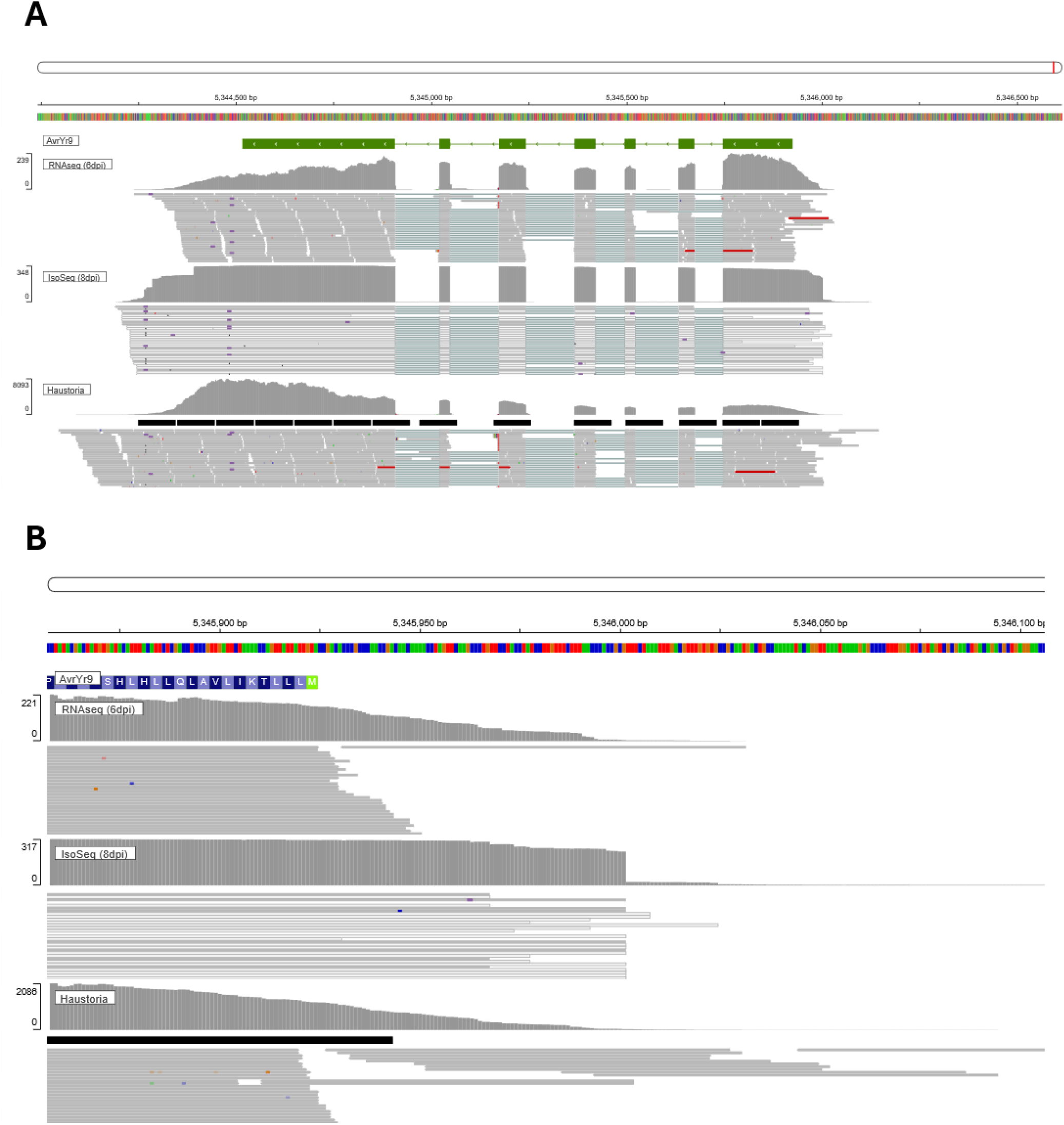
Transcript read mapping to *AvrYr9* gene in Pst104. The coding sequence of *AvrYr9* is shown as the top track.

**Supplementary Figure S6:**
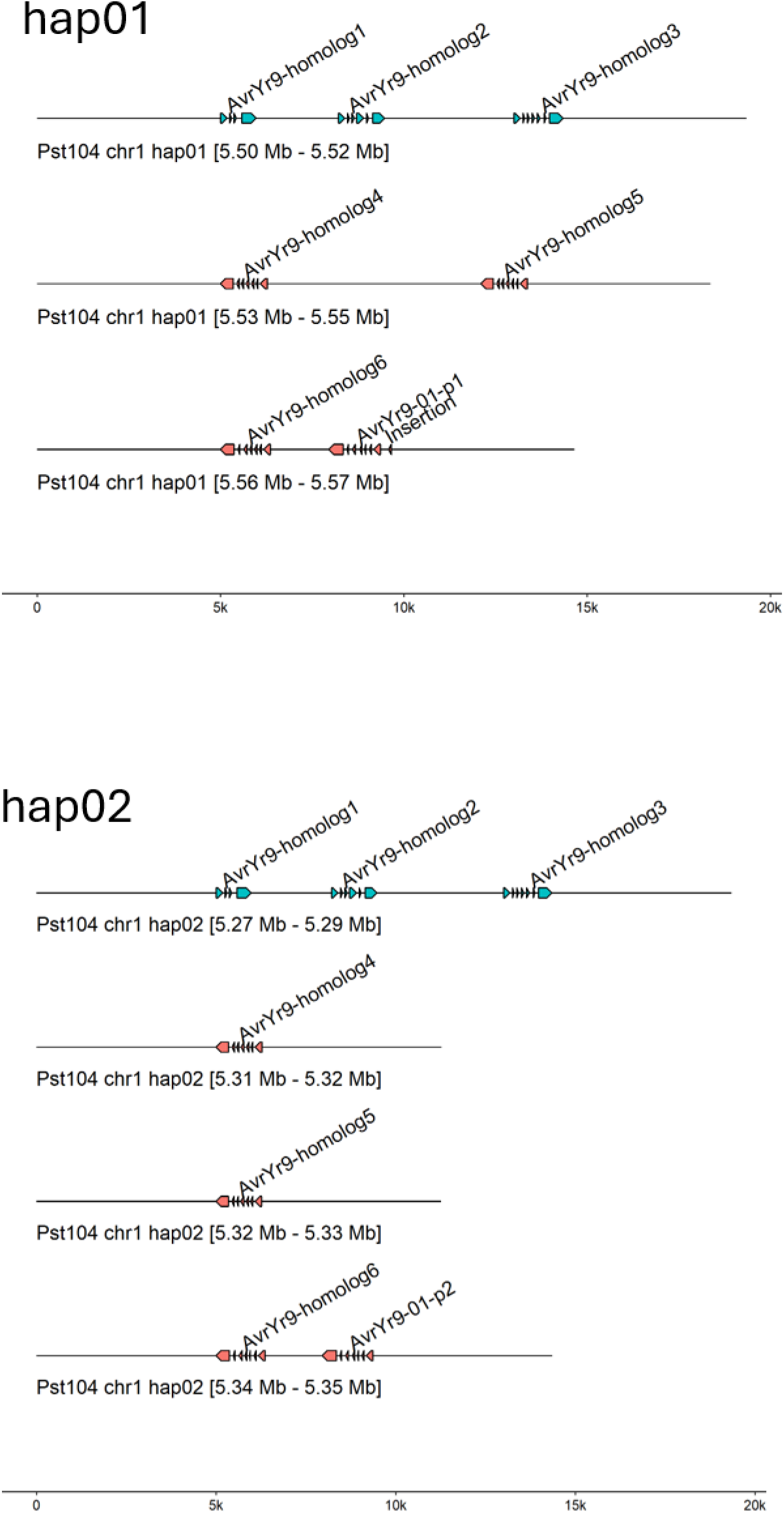
Structure of *AvrYr9* locus on chromosome 2 of Pst104 hap01 and hap02 showing arrangement of six related *AvrYr9*-homologous genes in a cluster on each haplotype.

**Supplementary Figure S7:**
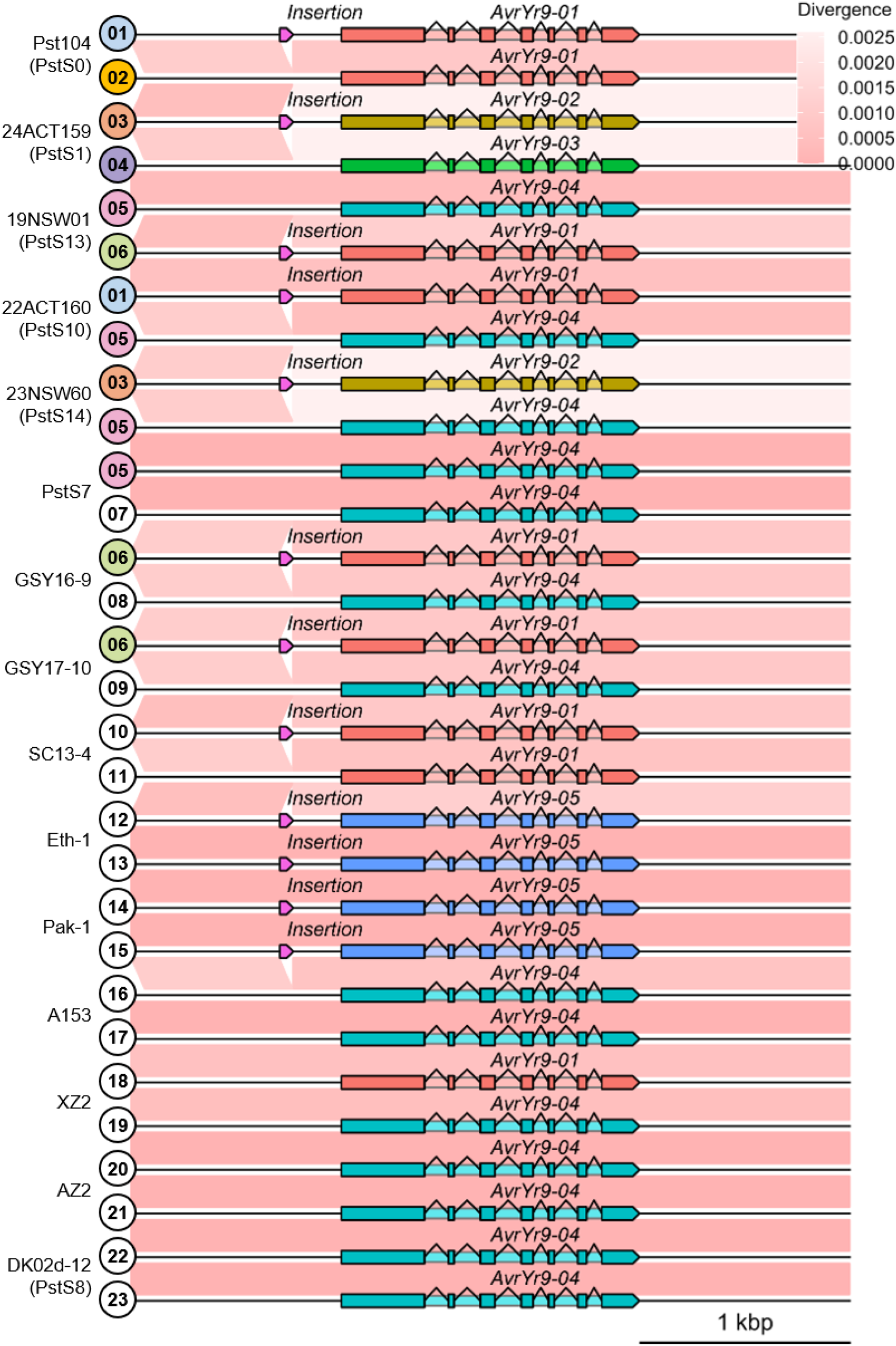
Synteny plot of the *AvrYr9* gene region. *AvrYr9* locus genotypes in haplotypes from fifteen *Pst* isolate genomes. Gene colors represent different sequence variants with pink shading indicating sequence similarity between haplotypes as per the scale inset. An insertion in the promoter region of some *AvrYr9* alleles is also indicated. *X*-axis shows the scale in kbp.

**Supplementary Figure S8:**
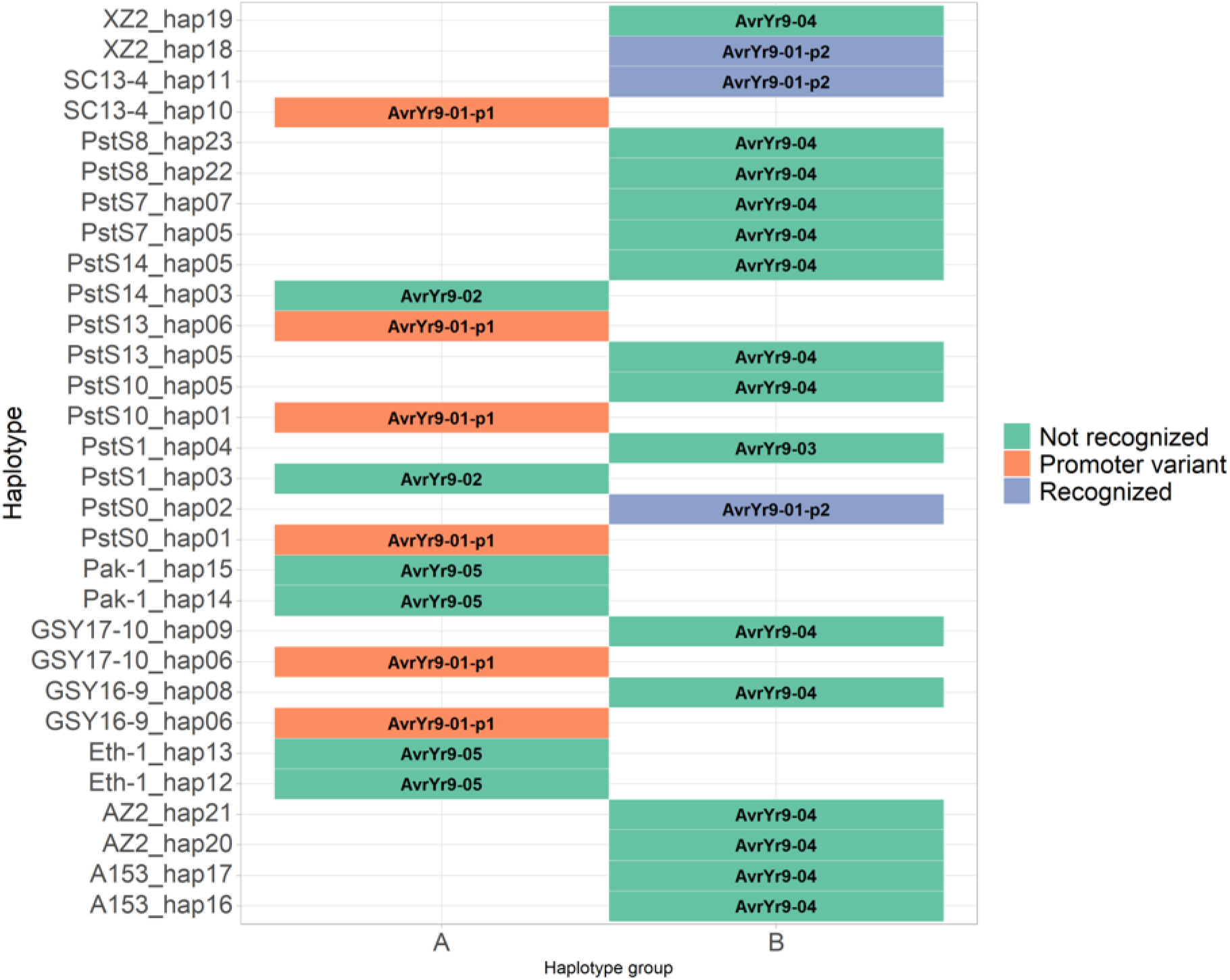
The *AvrYr9* locus on the Pst104 haplotypes. *Pst* genome assemblies are shown with presence of the *AvrYr9* variants. Haplotype groups are labelled according to A/B nomenclature (Wang *et al*., 2026a).

